# Dying oligodendrocytes persist without mitochondria

**DOI:** 10.64898/2026.01.19.699752

**Authors:** Xhoela Bame, S. Zoela Gilani, Yasmine Kamen, Robert A. Hill

## Abstract

Myelin is an insulating, multi-layered membrane that supports axonal integrity and neural communication. Different stressors impair myelinating oligodendrocytes, leading to demyelination, inflammation, and neurodegeneration. The intracellular processes underlying oligodendrocyte degeneration and death are unclear. Here, using optically targeted DNA damage that causes single-cell demyelination, we reveal that injured mature oligodendrocytes lose mitochondria within days and persist without them for weeks to months before cell death. This differs from other oligodendrocyte lineage cells, which exhibit acute mitochondrial changes followed by rapid cell death. Conditional deletion of the mitochondrial-related gene, *Fis1*, in mature oligodendrocytes, similarly causes acute loss of mitochondria and prolonged cell death. The unique cell death is characterized by nuclear changes, intracellular stress, and markers of disease-associated oligodendrocytes. Thus, mitochondrial loss may be an early marker of oligodendrocyte pathology, and mitochondrial quality control is required for oligodendrocyte and myelin homeostasis.

## INTRODUCTION

Myelin is a specialized structure that has evolved to insulate axonal tracts and enable efficient nerve conduction^1^. Myelin roles also extend to supporting neuronal metabolism and plasticity^2–4^, making it an important contributor to overall neural circuit health. New myelin is generated throughout life as oligodendrocyte precursor cells (OPCs) differentiate and transform into mature, myelinating oligodendrocytes^4,5^. Once oligodendrocytes and their myelin form, they are stable and persist for life in a healthy, adult brain^6–8^. However, as long-lived and post-mitotic cells, oligodendrocytes accumulate cellular stress throughout life, including oxidative stress and DNA damage, which can contribute to their degeneration^9–13^. Oligodendrocyte degeneration underlies the pathophysiology of multiple neurodegenerative diseases and aging, leading to impaired neural transmission, chronic neuroinflammation, and cognitive decline^6,14–18^.

Despite accumulating evidence for their involvement in pathology, the mechanisms driving their degeneration and death are unclear. Given the central role of mitochondria in multiple cell death pathways^19,20^, mitochondrial alterations are likely involved in oligodendrocyte death. In support of this, mitochondrial dysfunction and metabolic stress are associated with demyelinating conditions^12,21–29^, and can precede neurodegeneration and neuroinflammation^30–32^. In addition, mitochondrial dysfunction can impact the viability of the oligodendrocyte lineage cells^33–40^, and mitochondrial manipulations can restore oligodendrocyte loss upon demyelination^41,42^.

Mitochondrial homeostasis relies on tightly regulated processes of mitochondrial fission, fusion, transport, and turnover^43^. Different molecular regulators of these processes show differential expression throughout the oligodendrocyte lineage^33,36,44^, but their precise role and the stage-specific effects of their disruption in this lineage are unknown. One of these is *Fis1* (fission, mitochondrial 1), which encodes a mitochondrial outer membrane protein, FIS1 (mitochondrial fission 1 protein)^45^, and is upregulated upon oligodendrocyte maturation^33,44^. FIS1 is important for mitochondrial maintenance and quality control, spanning multiple functions across autophagy and its specialized form, mitophagy; mitochondrial dynamics and energetics; peroxisomal dynamics; and mitochondrial lysosomal recruitment^46–56^. Through these multifunctional roles, FIS1 impacts cell proliferation, maturation, and death in a context-dependent manner^50,54,57–59^. However, the role of FIS1 in the fate of oligodendrocyte lineage cells has not been explored.

Here, we caused DNA damage in single cells of the oligodendrocyte lineage and performed longitudinal intravital imaging to follow mitochondrial changes during their death. In addition, we determined how the conditional loss of *Fis1* in oligodendrocyte lineage cells affects their fate and maintenance. Our results reveal an acute loss of mitochondria during the prolonged death of mature oligodendrocytes, which is conserved across both DNA damage-based single-cell death and genetic deletion of *Fis1* in mature oligodendrocytes.

## RESULTS

### Different cell death dynamics across the oligodendrocyte lineage

To determine the spatiotemporal dynamics of mitochondria during the death of single oligodendrocyte lineage cells in the intact brain, we first generated a mouse line (*Cspg4*-creER; Ai9/PhAM) with expression of fluorescent reporters, tdTomato and mitochondrially targeted Dendra2 (mito-Dendra2), in the oligodendrocyte lineage. We performed cranial windows to visualize these cells in their native environment and used morphological and mitochondrial features^60^ to characterize their stage (Fig. 1, a and b, see Methods). The nuclei of the cells were labeled by a cell-permeable, DNA-binding dye, Hoechst 33342, during the surgical preparation over the somatosensory cortex. A two-photon femtosecond pulsed laser was used to irradiate a defined region over the nuclei of the targeted cells using a method called two-photon chemical apoptotic targeted ablation (2Phatal)^61^, resulting in photobleaching of the DNA-binding dye while keeping the cell intact (Fig. 1a). This procedure leads to potential reactive oxygen species-mediated DNA damage and programmed cell death^61^.

**Figure 1.**
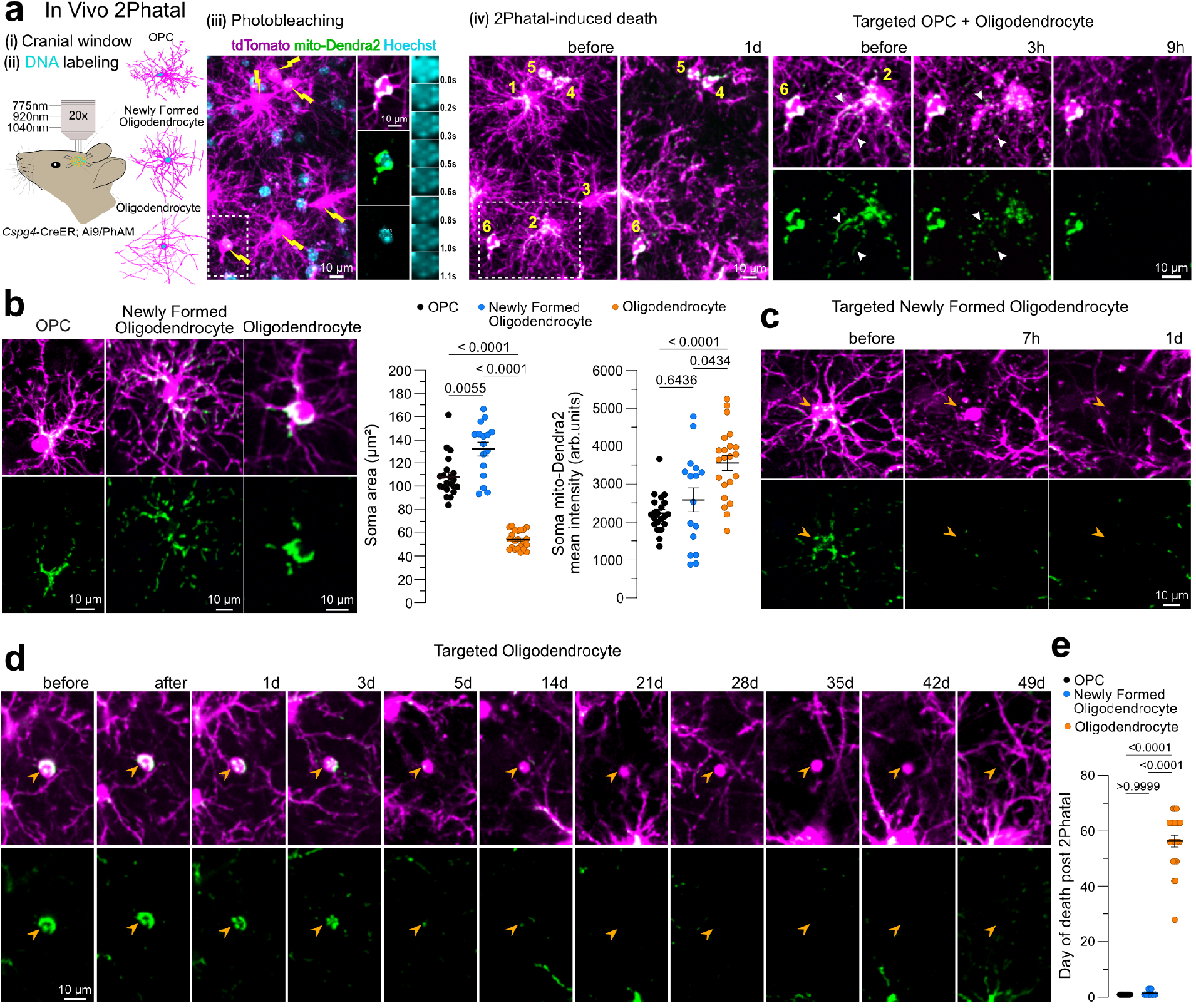
Temporal dynamics of cell death vary across the oligodendrocyte lineage. **a**, Schematic for inducing 2Phatal in vivo. 2Phatal-targeted OPCs (#1-3) were cleared within a day, whereas targeted oligodendrocytes (#4-6) persisted longer. White arrowheads point at fragmenting mitochondria in a dying OPC. **b**, Examples of distinct stages of the oligodendrocyte lineage and quantification of soma area and soma mitochondrial content (*n* = 21 OPCs, 16 newly formed oligodendrocytes, and 22 mature oligodendrocytes from 3-5 mice, Welch ANOVA with Dunnett’s T3 multiple comparisons test). **c**, Example of a 2Phatal-targeted newly formed oligodendrocyte that was cleared by the next day. **d**, Example of a 2Phatal-targeted mature oligodendrocyte persisting for over 6 weeks. Yellow arrowheads in **(c)** and **(d)**, point at targeted cells. **e**, Timing of cell soma clearance after 2-Phatal (*n* = 21 OPCs, 16 newly formed oligodendrocytes, and 22 mature oligodendrocytes from 3-5 mice, Kruskal Wallis with Dunn’s T3 multiple comparisons test). Data are shown as mean ± SEM.

Targeted OPCs underwent a stereotyped death process that involved fragmentation of mitochondria and formation of apoptotic bodies that were cleared in less than 24 hours post 2Phatal (Fig. 1a and fig. S1a). This mirrored the spontaneous death process that these cells undergo in the live brain (fig. S1b), potentially during their attempt to differentiate^60^, showing the high reliability of 2Phatal for mimicking physiological death. Targeted newly formed oligodendrocytes underwent a similar process, stretched over 1-3 days^62^ (Fig. 1c, and fig. S1, c and d). Targeted mature oligodendrocytes, on the other hand, despite similarly displaying mitochondrial fragmentation early on, underwent a more prolonged death process, lasting ∼1-2 months, highlighting the different cell death dynamics across the stages^62–64^ (Fig. 1, d and e, and fig. S1, e to g). Thus, single-cell 2Phatal coupled with endogenous fluorescent mitochondrial labeling allowed for the investigation of mitochondrial dynamics during the differing death dynamics of cells across the oligodendrocyte lineage.

### Mitochondria are rapidly lost during the slow oligodendrocyte death

Following 2Phatal-induced single-cell DNA damage, targeted mature oligodendrocytes were longitudinally imaged in vivo up to their clearance, falling between 4 and 10 weeks after the insult. Mitochondrial changes that accompanied the death of single oligodendrocytes were quantified and compared to those of untargeted cells within the same site, which served as internal controls (Fig. 2a). Mitochondrial shape analysis revealed an increase in mitochondrial circularity for the targeted cells within the first 3 days post 2Phatal (Fig. 2b), reflecting the early start of mitochondrial fragmentation, likely preceding mitochondrial degradation^65^. Fluorescence intensity quantifications showed an early and consistent decline in tdTomato signal within the targeted oligodendrocytes soma (Fig. 2c), attributed to the soma shrinkage, characteristic of certain forms of cell death^66^. Mito-Dendra2 signal within the oligodendrocytes somas also sharply declined within the first few days after 2Phatal (starting at day 3 in our analysis), with intensity measurements dropping progressively until the signal became minimal or undetectable, and remained such throughout the course of oligodendrocytes’ prolonged death (Fig. 2d). This decline was not solely due to the soma shrinkage, as the ratio of mito-Dendra2 to tdTomato signal, normalized to baseline, also declined over time for the targeted cells (Fig. 2e), indicating a disproportional loss of mito-Dendra2 signal compared to tdTomato. Additionally, this was unlikely to be caused by the repeated imaging, as both mito-Dendra2 and tdTomato signals remained stable in control cells that were longitudinally imaged in the same field of view (Fig. 2, a to e). Therefore, death-targeted oligodendrocytes quickly lost their mitochondria and persisted without them for several weeks, occasionally up to 2 months, until their clearance (Fig. 2, and fig. S1, e to g). Some oligodendrocytes maintained a few mito-Dendra2 puncta for longer (Fig. 2, e (cell#2) and f), however, their low number suggests limited capacity to support the metabolic needs of the cell, and their stable positioning over time further suggests limited turnover.

**Figure 2.**
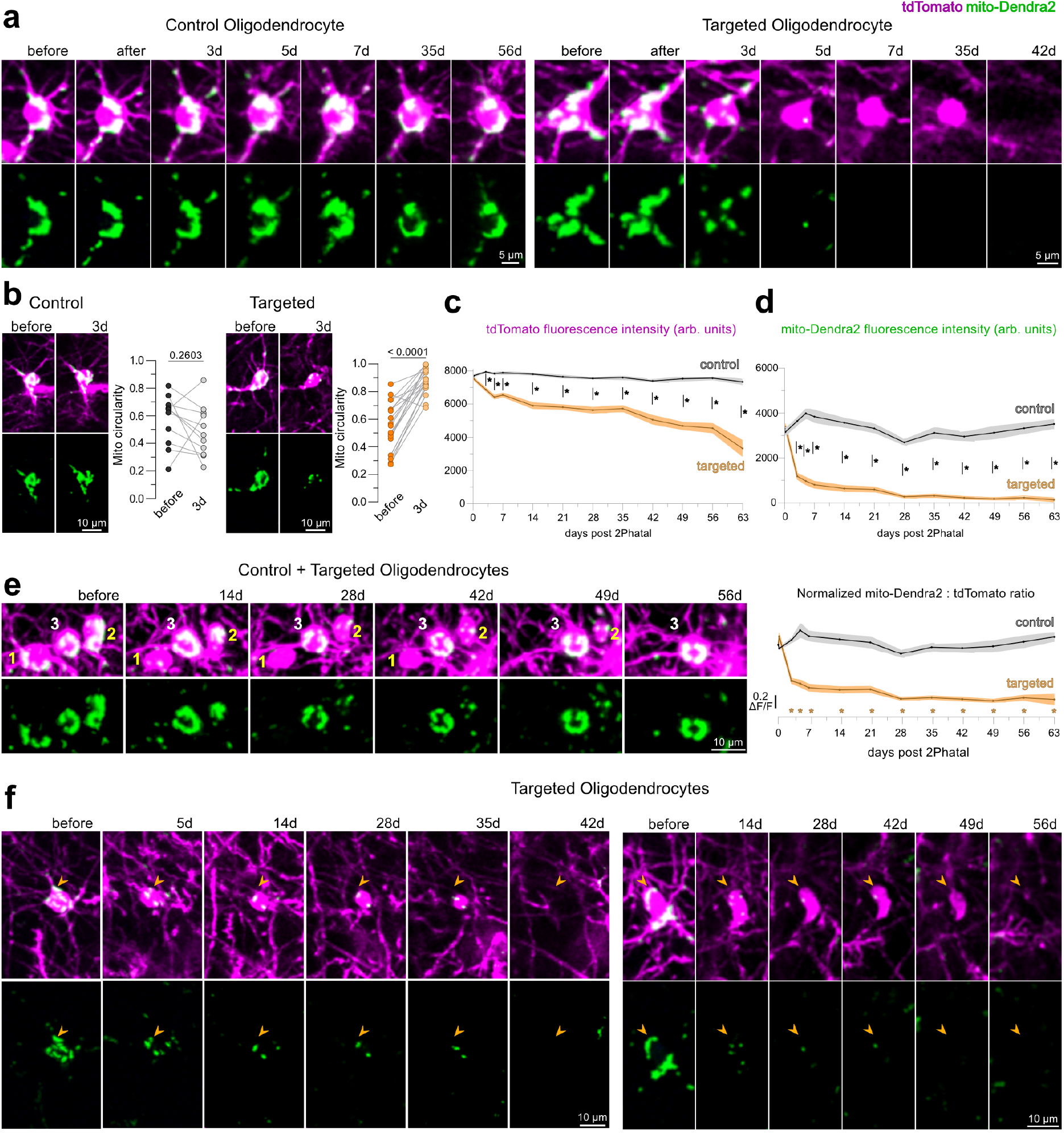
Rapid mitochondrial loss during slow oligodendrocyte death. **a**, Longitudinal imaging of a control (left) and a 2Phatal-targeted (right) oligodendrocyte. Note the targeted cell persists for ∼1 month after losing its mitochondria within 1 week after the injury. **b**, Images of a control and targeted oligodendrocytes before and 3 days after 2Phatal together with quantifications of mitochondrial circularity at each time-point (*n* = 11 control and 21 targeted oligodendrocytes from 3 mice, paired two-tailed t-test). **c, d**, Quantification of mean tdTomato **(c)** and mean mito-Dendra2 **(d)** fluorescence intensity within the soma of control and targeted oligodendrocytes (*n* = 11 control and 22 targeted oligodendrocytes from 3 mice, multiple unpaired two-sample t-tests with the Holm-Šídák method). Asterisks indicate statistical significance at <0.000001. **e**, Longitudinal images of targeted (yellow numbered, #1 and #2) and control (white numbered, #3) oligodendrocytes. Quantification of mito-Dendra2 to tdTomato fluorescence intensity normalized to the time point before 2Phatal. Asterisks indicate data points that significantly deviate from 0 at the 99% confidence interval, corresponding to the targeted cells group (*n* = 11 control and 22 targeted oligodendrocytes from 3 mice). **f**, Longitudinal images of targeted oligodendrocytes that lost most of their mitochondrial signal early on but maintained a few mitochondrial puncta up until or near their clearance. Arrowheads point at the targeted cells. In graphs **c-e**, data were analyzed up to the last timepoint preceding cell clearance. Data are shown as mean ± SEM.

To confirm that the observed mito-Dendra2 loss was not a technical artifact, but rather a bona fide response to the injury, we validated the results using an independent marker, TOM20, a stable, outer mitochondrial membrane protein. We first scaled up the 2Phatal technique and used it to target around 500 oligodendrocytes per mouse in vivo (hereafter called multifocal 2Phatal). Twenty-eight days after the multifocal 2Phatal, a time point when, as established by intravital imaging (Fig. 1, Fig. 2, and Fig. 3a), oligodendrocytes were still present, but had lost most of their Dendra2-labeled mitochondria, the mice were perfused, and brains were collected in order to stain for additional markers in fixed tissue (mitochondrial marker TOM20, oligodendrocyte marker CNP, and nucleus marker Hoechst 33342). Fixed tissue imaging was done in the somatosensory cortex, in regions corresponding to the site of multifocal 2Phatal. Targeted cells were identified based on their condensed soma and nucleus (Fig. 3, b to e), typical hallmarks of cell death^66^. Their nuclei also displayed aberrant morphology, and nucleoli were no longer discernible (Fig. 3, b, d, and e). Quantification of mito-Dendra2 and TOM20 fluorescence intensity confirmed the intravital imaging results, showing that both mitochondrial signals significantly declined in targeted cells compared to control cells (Fig. 3c). In some cases, no overlap was seen between remaining mito-Dendra2 signal in targeted cells and TOM20 labeling (Fig. 3, c and d), suggesting compromised mitochondrial integrity and potential structural damage or delayed degradation of the mito-Dendra2 fluorophore. Overall, these results show that dying oligodendrocytes have few to no mitochondria left well before their death and clearance. This suggests a potential metabolic adaptation and a distinct mode of death for these cells in which they persist for weeks to months despite mitochondrial loss, unlike the other cells in their lineage that get cleared within hours to days following comparable levels of injury and mitochondrial shape alterations.

**Figure 3.**
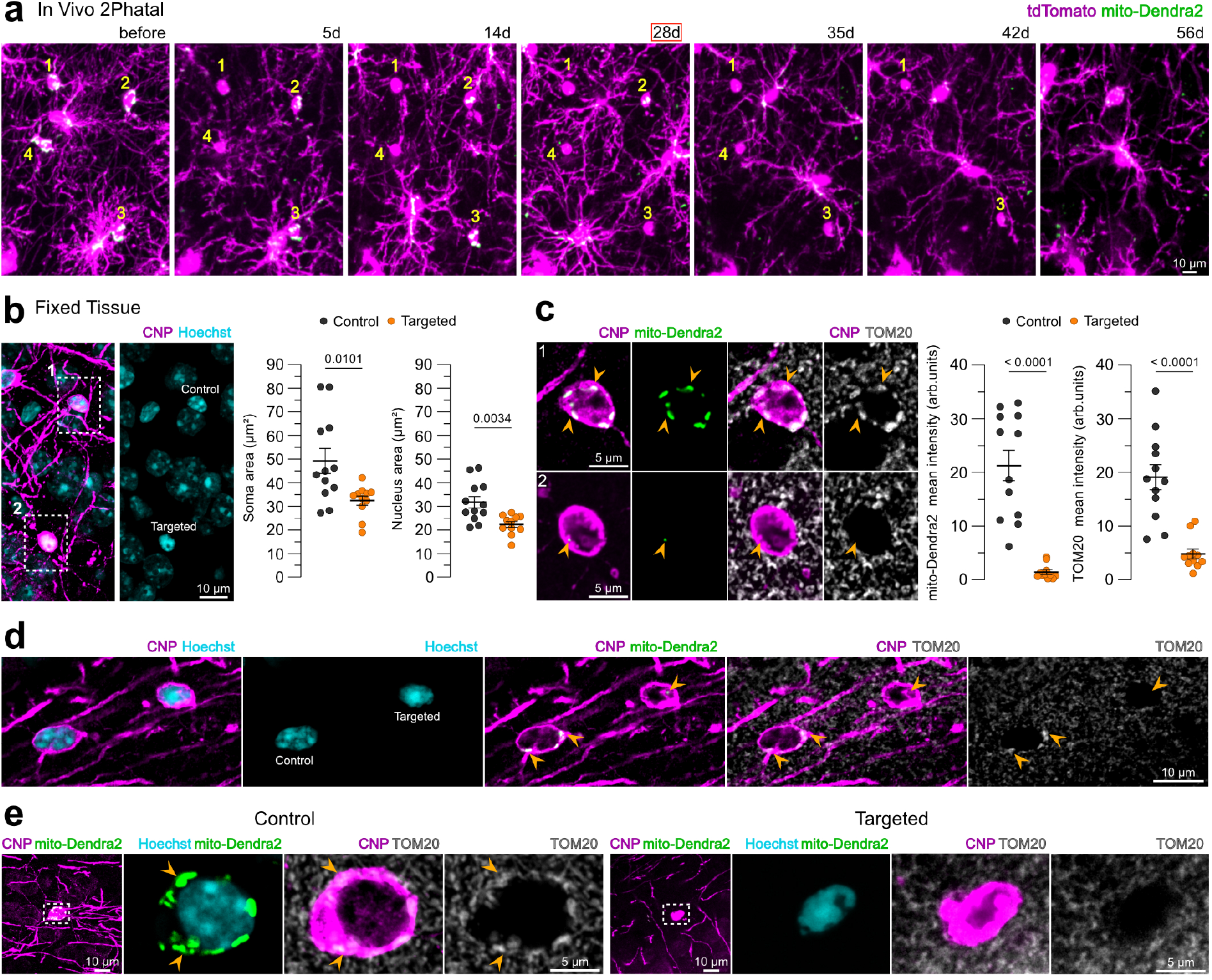
Dying oligodendrocytes lack markers of mitochondria. **a**, Longitudinal imaging of 2Phatal-targeted oligodendrocytes (#1-4). Note that all cells were present on day 28, with most of their mitochondria lost. Oligodendrocyte clearance occurred by day 35 for #2; day 42 for #4, and day 56 for #1 and #3 following the injury. **b**, Cortical oligodendrocytes in fixed tissue at 28 days post injury. Quantification of soma and nucleus area. **c**, Zoomed images from single optical sections of cells in **(b)**, stained for TOM20. Quantification of soma mito-Dendra2 and TOM20 mean fluorescence intensity. In **(b)** and **(c)**, *n* = 12 control and 11 targeted oligodendrocytes from 3 mice, unpaired two-tailed *t* test with Welch’s correction for unequal variance. **d, e** Representative images of single optical sections showing control and targeted oligodendrocytes in fixed tissue. Note the lack of TOM20 immunoreactivity in the targeted cells, despite the remaining mito-Dendra2 punctum in **(d)**. Arrowheads point at mito-Dendra2 labeling. Data are shown as mean ± SEM.

### FIS1 is abundant in oligodendrocytes, yet dispensable for their generation

Given the dynamic reorganization of mitochondria during the generation of oligodendrocytes^60^ and mitochondrial alterations happening early on during their death (Fig. 2), we next asked whether manipulating a mitochondrial-related gene that was differentially expressed across the oligodendrocyte lineage, could affect their fate and maintenance. One of these genes, *Fis1*, is upregulated in cultured mature oligodendrocytes, at both transcript and protein levels, as shown by RNA assays and immunoblotting^36,33,44,67^. In addition, mature oligodendrocytes show the highest *Fis1* mRNA expression across all brain cells in both mice^44^ and humans^67^. We confirmed the corresponding protein-level changes in vivo by immunostaining for FIS1 in fixed brain tissue. Indeed, mature oligodendrocytes labeled by the CAII marker showed strong FIS1 immunostaining signal that distinguished them from the other neighboring cells in the tissue (Fig. 4a). Quantitative fluorescence intensity measurements showed greater FIS1 levels in mature oligodendrocytes compared to OPCs, marked by PDGFRA (Fig. 4b).

**Figure 4.**
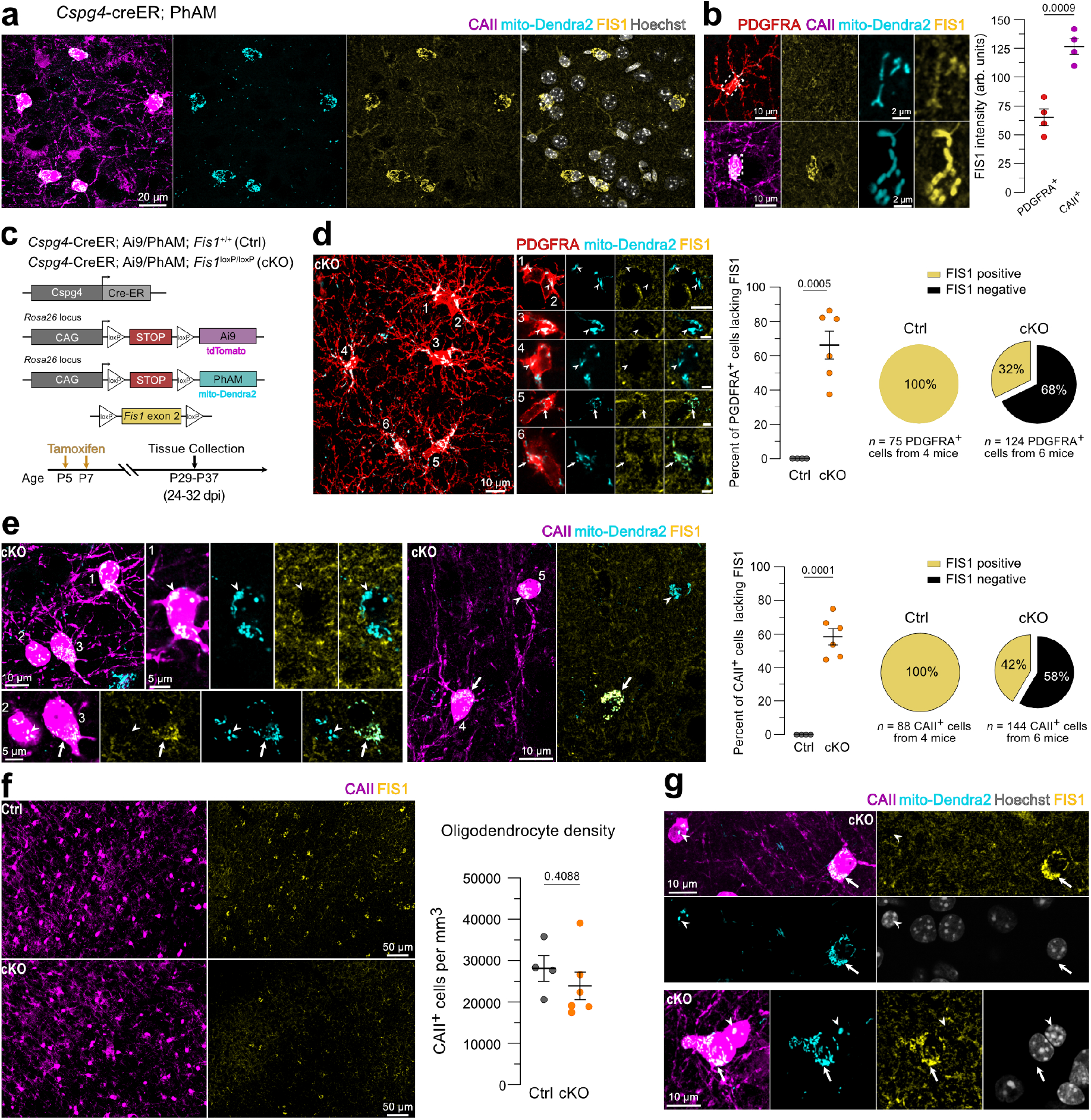
Oligodendrocyte generation is maintained in the absence of *Fis1*. **a**, FIS1 immunostaining in fixed tissue from the cerebral cortex of a *Cspg4*-CreER; PhAM transgenic mouse. Note that CAII-labeled oligodendrocytes show the strongest FIS1 signal and stand out from the surrounding cells. **b**, Representative images of a PDGFRA-labeled OPC (top) and CAII-labeled oligodendrocyte (bottom) along with their mito-Dendra2 and FIS1 signals. Boxes indicate regions shown at higher magnification in single optical z-sections. Quantification of FIS1 intensity within the mitochondrial ROIs from 23 PDGFRA^+^ and 23 CAII^+^ cells (*n* = 4 mice, unpaired two-tailed *t* test). **c**, Experimental timeline and genetic strategy for conditional deletion of *Fis1* and expression of mito-Dendra2 and tdTomato in *Cspg4* lineage cells. Two doses of tamoxifen were injected postnatally at P5 and P7 to induce Cre recombination, and experiments were performed 3-4 weeks later. **d**, Representative images of PDGFRA^+^ cells in the cKO tissue. A subset of the cells lacks FIS1 (#1-4), whereas the FIS1 signal is present in the others (#5-6). Scale bars in cropped images represent 2 µm. Percent of PDGFRA^+^ cells lacking FIS1 immunoreactivity in control and cKO tissue. **e**, Representative images of CAII^+^ cells in the cKO tissue. A subset of them lacks FIS1 (#1, #2, #5), whereas FIS1 signal is present in the others (#3, #4). Percent of CAII^+^ cells lacking FIS1 immunoreactivity in control and cKO tissue. In **d** and **e**, *n* = 4 control and 6 cKO mice, unpaired two-tailed *t* test with Welch’s correction for unequal variance. **f**, Representative fields of view from control and cKO tissue showing CAII and FIS1 immunostaining. Quantification of cortical oligodendrocyte density (*n* = 4 control and 6 cKO mice, unpaired two-tailed *t* test). **g**, Examples of FIS1-negative oligodendrocytes with disrupted mitochondria in the cKO tissue, along with adjacent FIS1-positive cells. In **d, e**, and **g**, arrowheads point at FIS1-negative cells, and arrows point at FIS1-positive cells and their mitochondria. dpi: days post tamoxifen injection. Data are shown as mean ± SEM.

Next, to specifically determine if *Fis1* plays a role in OPC fate or oligodendrocyte development, we generated a mouse line with conditional deletion of *Fis1* in OPCs and expression of tdTomato and mito-Dendra2 fluorescent reporters in these cells and their progeny (*Cspg4*-CreER; Ai9/PhAM; *Fis1*^loxP/loxP^). Tamoxifen was injected postnatally, at P5 and P7, to induce Cre-mediated recombination early during development, and prior to the onset of OPC differentiation. Experiments and tissue collection were performed 3-4 weeks later to be able to potentially capture early effects and minimize compensatory mechanisms that may occur at later stages (Fig. 4c). To confirm the *Fis1* deletion efficiency, we stained for FIS1 and assessed its presence in OPCs. Sixty-eight percent of PDGFRA-labeled OPCs lacked FIS1 immunoreactivity, reflective of a successful deletion of the gene in the majority of cells (Fig. 4d). Next, we assessed whether *Fis1* loss led to changes in mitochondrial morphometrics in OPCs. Comparisons were made within the *Fis1* conditional knockout (cKO) tissue between the FIS1-negative and FIS1-positive OPCs (fig. S2a), which served as internal controls to account for potential fixation artifacts that could alter mitochondria in the tissue^68^. Neither the content, size, nor morphology of mitochondria showed a difference between the two populations (fig. S2b). *Fis1* loss also did not affect OPC branching morphology as quantified by Sholl analysis (fig. S2c). We next tested for any changes in their proliferation or survival. OPC proliferation rates were comparable between the littermate control and cKO groups, as shown by EdU incorporation and KI67 immunostaining (fig. S3, a and b). Moreover, no overlap was seen between PDGFRA-positive cells and the programmed cell death marker, cleaved caspase-3 (CC3), in control or cKO tissue (fig. S3c). In addition, the density of apoptotic bodies derived from tdTomato-labeled cells and detected by intravital imaging also remained unchanged (fig. S3d), altogether indicating no detectable differences in death rate. The total density of EdU (fig. S3a) and CC3-labeled cells (fig. S3c) across the cerebral cortex was not different between control and cKO tissue, suggesting no off-target effects within the brain.

Given the high levels of FIS1 in mature oligodendrocytes, we next determined whether FIS1 regulates oligodendrocyte generation. We first checked for the presence of oligodendrocytes lacking FIS1 in the cKO tissue, which would infer that they were derived from OPCs with successful *Fis1* deletion, as Cre recombination was induced at the OPC stage and would be retained after they transition into mature cells. Indeed, 58% of mature oligodendrocytes lacked FIS1 (Fig. 4e), supporting their successful differentiation from recombined OPCs, while the rest were likely derived from OPCs that escaped the *Fis1* deletion. This value is consistent with the recombination percentage in OPCs (Fig. 4d), and together with the unchanged oligodendrocyte density between control and cKO groups at ∼1 month after recombination (Fig. 4f), suggests that oligodendrocyte generation was not impaired. Interestingly, low levels of mitochondria were observed in a small number of the mature oligodendrocytes that lacked FIS1. This was observed only in the cKO groups but never in the controls (Fig. 4g). These data hinted that the maintenance and homeostasis of the oligodendrocyte lineage could be affected in the long-term and prompted additional experiments and later time points in a new cohort of mice.

### Disruption of *Fis1* in the oligodendrocyte lineage leads to progressive loss of oligodendrocytes and their myelin

To observe if any effect arising from *Fis1* deletion in the oligodendrocyte lineage became apparent at a later stage, especially given the stable nature of the oligodendrocytes, we analyzed tissue from a new cohort of animals 3 months after Cre-mediated deletion of *Fis1* (Fig. 5a). Consistent with the 1-month cohort, we occasionally saw FIS1-negative oligodendrocytes marked by CAII showing hallmarks of cell death, including missing mitochondrial labeling and altered nuclear morphology (Fig. 5b), hinting at degeneration of this population. To test this, we first quantified the number of oligodendrocytes lacking FIS1 in the cKO tissue. This population comprised 20% of all mature oligodendrocytes (Fig. 5c), corresponding to a decline from the 58% at the 1-month time-point (Fig. 4e). Additionally, the density of oligodendrocytes labeled independently by CAII, ASPA, and CNP, showed a consistent decline by, on average, more than a third compared with littermate controls (Fig. 5, c and d). Myelin basic protein (MBP) labeling was also significantly lower in the cKO tissue compared to the control (Fig. 5d), altogether suggesting that over time, oligodendrocytes were depleted, despite their initial successful generation following *Fis1* deletion in OPCs (Fig. 4). To observe the response to demyelination and potential remyelination attempts, we stained for BCAS1, a marker of differentiating oligodendrocytes^69^. While the total density of BCAS1^+^ cells remained unchanged, there appeared to be more BCAS1-labeled cells with myelinating morphology in the cKO compared to the control. To quantify this, the overlap between BCAS1 and MBP was measured, revealing significantly more overlap in the cKO compared to the control (Fig. 5e). These data point toward more new myelin present in the cKO brain, likely indicating remyelination occurring from differentiating OPCs. While there were changes in oligodendrocytes and myelin, the OPC density was not different between the control and cKO (Fig. 5f). Additionally, FIS1-negative oligodendrocytes declined to 1% of all quantified CAII^+^ cells at 1-year post-recombination, supporting a gradual replacement of this population (fig. S4). Overall, these data suggest a progressive decline of FIS1-negative mature oligodendrocytes, which are slowly replaced by OPCs that escaped the *Fis1* deletion.

**Figure 5.**
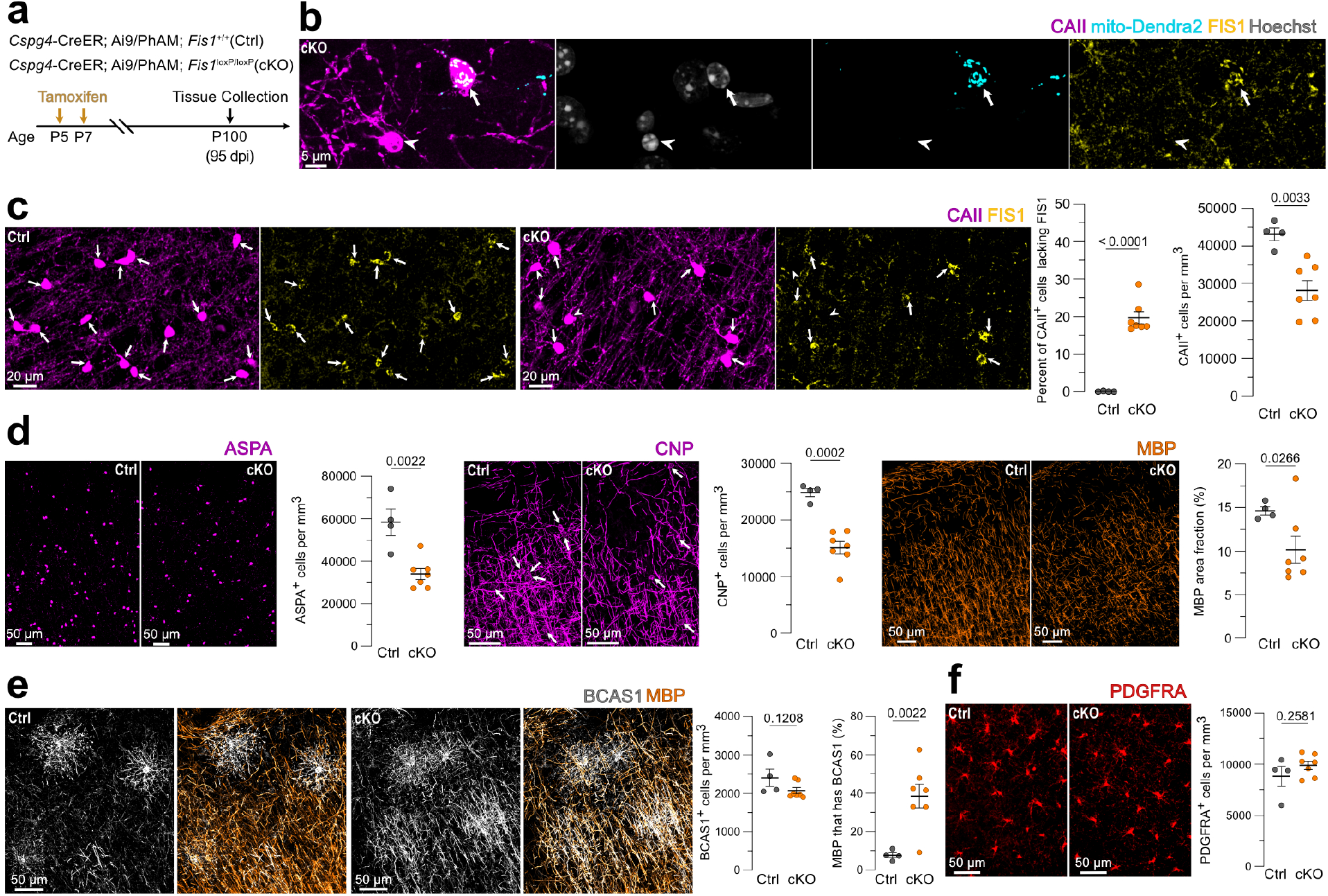
Delayed loss of oligodendrocytes and myelin after *Fis1* deletion in OPCs. **a**, Experimental timeline showing tamoxifen injections at P5 and P7 and tissue collection at P100. **b**, Example of a FIS1-positive oligodendrocyte adjacent to a FIS1-negative oligodendrocyte. Note the condensed nucleus and lost mitochondria in the latter. **c**, Representative images of CA-II labeled oligodendrocytes and FIS1 immunostaining in control and cKO tissue. Quantification of the percentage of CAII^+^ cells lacking FIS1 immunoreactivity and CAII^+^ cell density (*n* = 4 control and 7 cKO mice, unpaired two-tailed *t* test; Welch’s correction for unequal variance was applied for the first graph, left to right). In **(b)** and **(c)**, arrows point at FIS1-positive oligodendrocytes, and arrowheads point at FIS1-negative oligodendrocytes. **d**, Representative images of ASPA, CNP, and MBP staining across control and cKO groups, along with quantification of cell densities and MBP area coverage (*n* = 4 control and 7 cKO mice, unpaired two-tailed *t* test; Welch’s correction was applied for the MBP graph). Arrows point at CNP^+^ cells. **e**, Representative images of BCAS1 and MBP staining, along with BCAS1^+^ cell density and BCAS1 signal coverage within MBP^+^ structures across the two groups (*n* = 4 control and 7 cKO mice, unpaired two-tailed *t* test; Welch’s correction was applied for the second graph, left to right). **f**, Representative images of PDGFRA staining along with PDGFRA^+^ cell densities in control and cKO mice (*n* = 4 control and 7 cKO mice, unpaired two-tailed *t* test). Data are shown as mean ± SEM.

Of note, the FIS1-negative OPC population was also mostly replaced at the 3-month time point, with only 2% of the cells remaining (5/245 PDGFRA^+^ cells from 4 cKO mice), suggesting potential missed effects of the deletion in this population, which, likely due to their ability to reach population homeostasis quickly^70–72^, was not apparent in our analysis.

### Disruption of *Fis1* in mature oligodendrocytes results in loss of mitochondria and hallmarks of slow oligodendrocyte death

Given the high baseline levels of FIS1 in mature oligodendrocytes, and their progressive loss following conditional deletion of *Fis1* in their precursors, we next tested if this represented an oligodendrocyte-specific effect, and what early defects could contribute to it. Thus, we generated another line, *Plp1*-CreER; PhAM; *Fis1*^loxP/loxP^, with conditional deletion of *Fis1* and expression of mito-Dendra2 specifically in myelinating cells, including more mature oligodendrocytes. Tamoxifen administration started at 2 months of age, after peak developmental myelination, to predominantly induce recombination in fully mature oligodendrocytes. Injections were spread throughout 6 weeks to ensure sustained Cre recombination of newly generated oligodendrocytes and tissue was collected 2 months after the first injection (Fig. 6a). This resulted in a mixed population of oligodendrocytes with early and late *Fis1* loss at the start of the experiments, allowing for the detection of transient effects of the deletion and any potential phenotypes prior to any major depletion of the oligodendrocyte population. The recombination efficiency was high, as 93% of the oligodendrocytes in the cKO tissue lacked FIS1 (Fig. 6b). Within the FIS1-negative oligodendrocytes, those with an altered nucleus morphology and low mitochondrial content were seen more frequently than in examples from Figs. 4g and 5b. Given that this phenotype was more widespread in this tissue, we formally quantified the number of oligodendrocytes with mito-Dendra2 loss. Between 40-50% of the oligodendrocytes displayed either a low mito-Dendra2 signal or a complete lack of it, as quantified by both, experimentally blinded manual (Fig. 6c), and automated (fig. S5a) approaches. In addition, oligodendrocytes in the cKO tissue displayed a smaller nucleus and soma size, with the latter being evident in analysis by cell (fig. S5b) but not reaching statistical significance in our analysis by animal (Fig. 6d), likely due to the heterogeneity of the cells and stages. The decrease in mitochondrial content in the cKO group, was confirmed by quantifying fluorescence from both, the genetically encoded mito-Dendra2, and TOM20 immunostaining, showing comparable results (Fig. 6, e and f, and fig. S5b) and supporting the use of mito-Dendra2 as a reliable mitochondrial marker.

**Figure 6.**
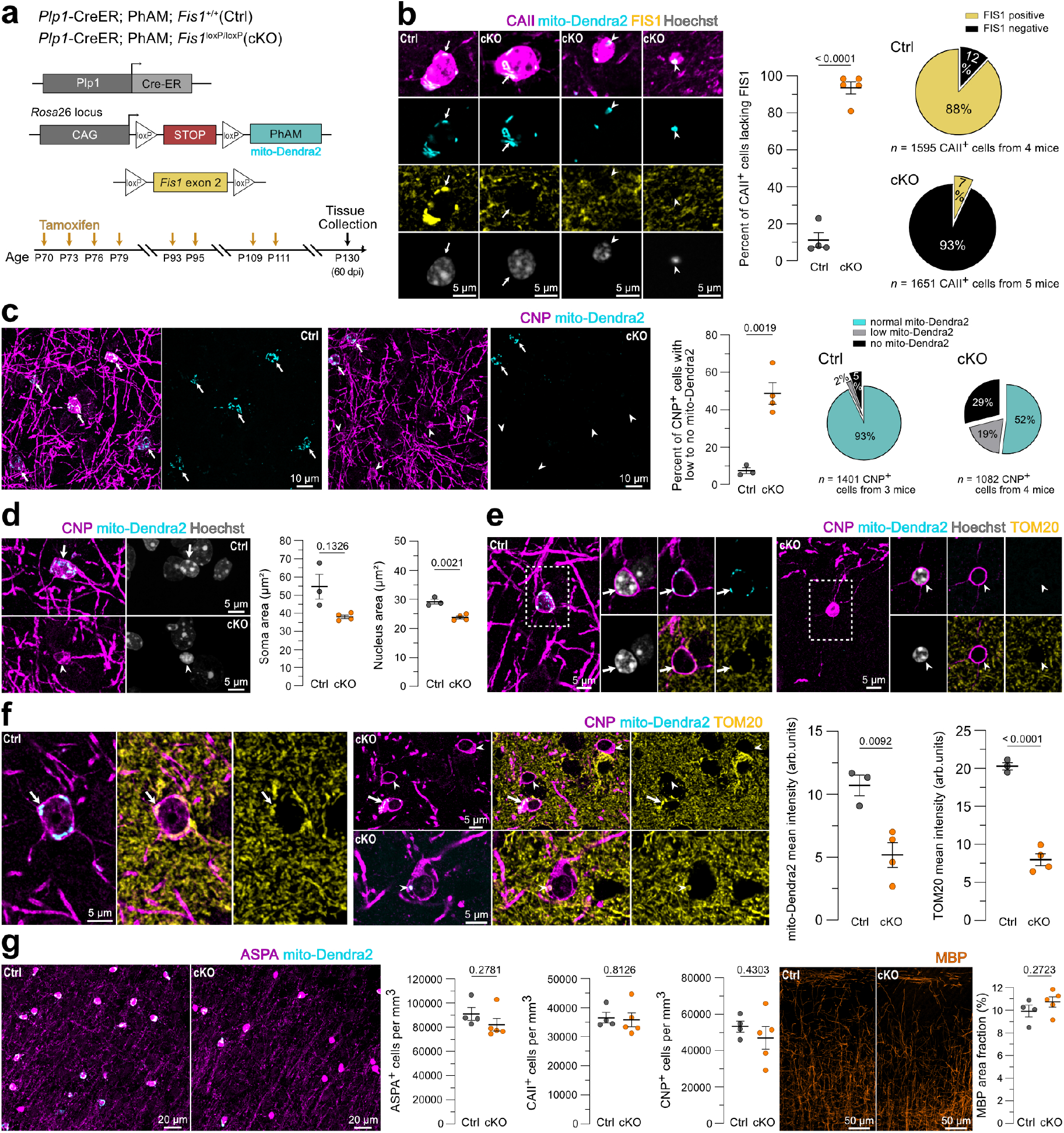
*Fis1* deletion in mature oligodendrocytes causes mitochondrial loss and nuclear condensation. **a**, Experimental timeline and genetic strategy for conditional deletion of *Fis1* and expression of mito-Dendra2 in myelinating cells. Eight doses of tamoxifen were administered starting at ∼P70, and experiments were performed 2 months after the first injection. **b**, Representative images showing oligodendrocytes in the control and cKO tissue, and heterogeneous phenotypes of FIS1-negative oligodendrocytes. Automated quantification of the percentage of CAII-labeled oligodendrocytes lacking FIS1 immunoreactivity (*n* = 4 control and 5 cKO mice, unpaired two-tailed *t* test). **c**, Representative images of CNP-labeled oligodendrocytes and their mitochondria across control and cKO groups. Note the lack of mito-Dendra2 signal in a subset of cells in the cKO tissue. Quantification of the percentage of CNP^+^ cells with low or no mito-Dendra2 signal (*n* = 3 control and 4 cKO mice, unpaired two-tailed *t* test). **d**, Zoomed in examples of oligodendrocytes and their nuclei from fields of view in **(c)**. Quantification of soma and nucleus area (*n* = 3 control and 4 cKO mice, unpaired two-tailed *t* test; Welch’s correction for unequal variance was applied for the first graph, left to right). **e, f**, Examples of CNP-labeled oligodendrocytes and TOM20 immunostaining. Note that TOM20 signal corresponds to the presence or absence of the mito-Dendra2 signal, with some contributions from surrounding cells in single optical sections. Quantification of mito-Dendra2 and TOM20 mean fluorescence intensity in the somas of CNP-labeled oligodendrocytes (*n* = 3 control and 4 cKO mice, unpaired two-tailed *t* test). In **b-f**, arrows point at cells with normal mito-Dendra2 signal, and arrowheads point at cells with low or no mito-Dendra2 signal. **g**, ASPA^+^, CAII^+^, and CNP^+^ cell density and MBP area coverage in control and cKO groups (*n* = 4 control and 5 cKO mice, unpaired two-tailed *t* test). Data are shown as mean ± SEM.

Overall, the smaller nuclei and the confirmed loss of mitochondria in oligodendrocytes in the cKO group, were consistent with the features displayed by oligodendrocytes targeted by 2Phatal (Fig. 3), and hallmarks of death, suggesting that they are likely undergoing degeneration. However, the density of oligodendrocytes and myelin coverage remained unchanged between the littermate control and cKO groups (Fig. 6g), likely due to the prolonged process of degeneration of these cells. The number of differentiating oligodendrocytes marked by procaspase-3 (pro-CASP3)^73^ was also not different between cKO and control groups (fig. S6), suggesting comparable differentiation attempts.

Next, we looked for the presence of cell death markers in these cells. While a similar number of cells stained positive for CC3 across the two groups (fig. S7a), no signal was detected in CAII-labeled oligodendrocytes, including those with condensed nuclei and/or mitochondrial loss (fig. S7b). This is consistent with a lack of CC3 and CAII dual-positive population in other models of demyelination^62^, suggesting a lack of caspase-dependent cell death in these cells. We also looked for indicators of parthanatos, a non-apoptotic form of cell death, characterized by a redistribution of poly (ADP-ribose) (PAR)^74^ and implicated in demyelinating pathologies^62,75^. However, PAR subcellular localization was similar across both groups, and primarily confined to the cell nucleus regardless of the mitochondrial levels in the cell (fig. S7, c to e), suggesting that parthanatos was unlikely to occur at this stage. In addition, we observed no indication of an inflammatory response from microglia (fig. S7f) or astrocytes (fig. S7g). Thus, despite showing hallmarks of death upon FIS1 loss, oligodendrocyte clearance likely occurs later, consistent with them being rapidly phagocytosed months after catastrophic DNA damage^76^.

### Oligodendrocytes lacking FIS1 show elevated levels of cell stress markers

With a loss of mitochondria and compromised nuclear morphology following *Fis1* deletion in oligodendrocytes, we next looked for autophagy and stress-related markers potentially linked to these changes. One of these was P62 (SQSTM1), a multifunctional protein and an autophagy adaptor shown to play a role in cellular stress responses and pathways regulating cell survival and death^77,78^. In addition, P62 is shown to accumulate upon *Fis1* loss in certain cellular contexts^54^ as well as in aging and autophagy-impaired cells of the oligodendrocyte lineage^79^, relating to both the health and maintenance of these cells and the *Fis1* loss consequences. Oligodendrocytes that maintained their mitochondria in the cKO tissue, likely due to having lost *Fis1* more recently, showed similar P62 levels to cells in control tissue (Fig. 7a). However, a mixture of high and low P62 intensity signals was seen in the oligodendrocytes that had mostly lost their mitochondria and were at later stages of degeneration (Fig. 7a). Importantly, high P62 levels were exclusively seen in this group, which may point to impaired autophagy and proteotoxic stress in these cells.

**Figure 7.**
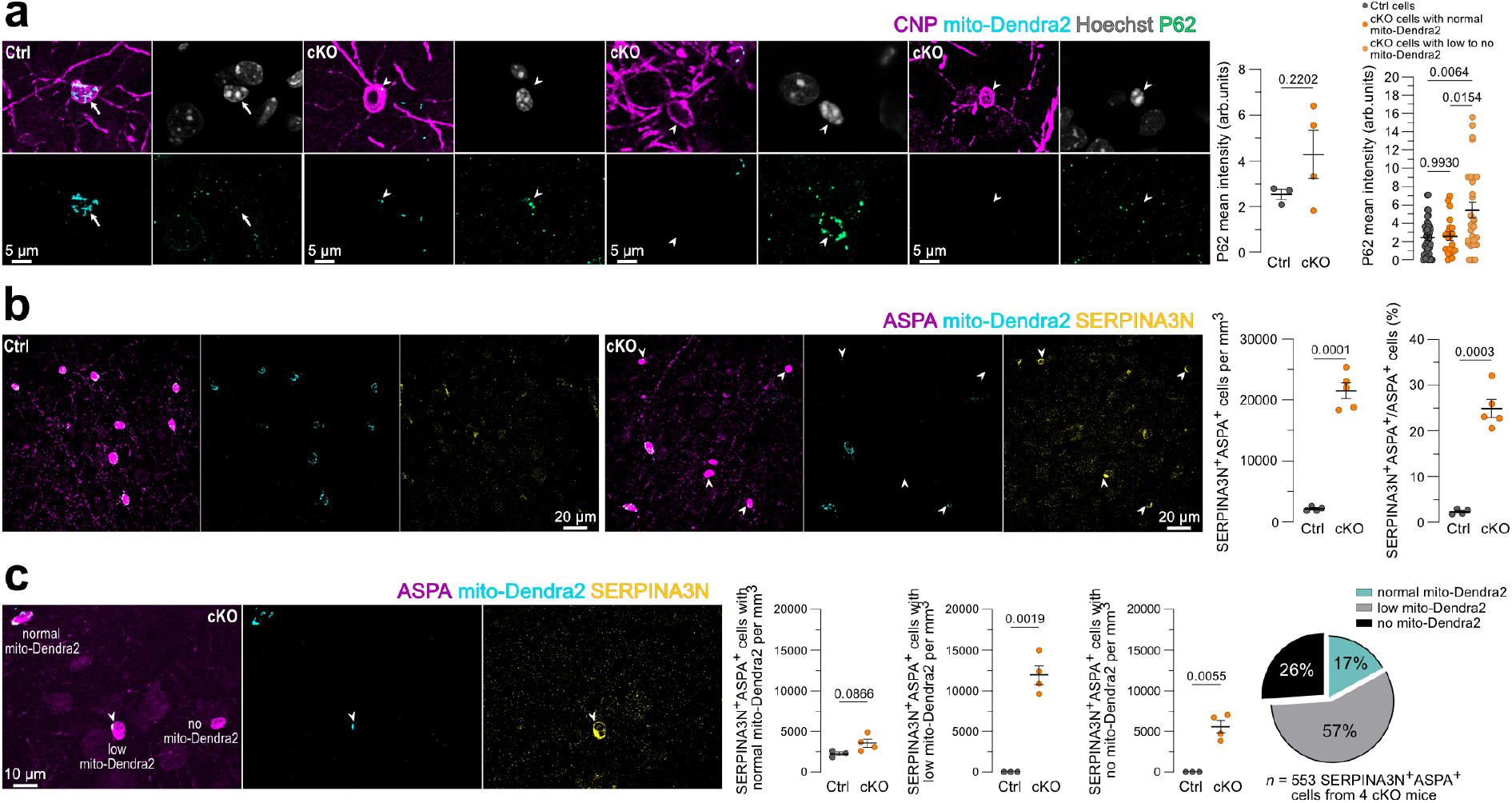
Loss of *Fis1* in mature oligodendrocytes leads to cell stress and increased SERPINA3N. **a**, Representative images of P62 immunostaining in oligodendrocytes from control and cKO, *Plp1*-CreER; PhAM; *Fis1*^loxP/loxP^ mice. Quantification of P62 mean fluorescence intensity per animal (*n* = 3 control and 4 cKO mice, unpaired two-tailed *t* test) and per cell (*n* = 39 cells from 3 control mice; 20 cells with normal mito-Dendra2, and 29 cells with low to no mito-Dendra2 from 4 cKO mice, Welch ANOVA with Dunnett’s T3 multiple comparisons test). Arrows point at CNP^+^ cells with normal mitochondria, and arrowheads point at CNP^+^ cells with low or no mito-Dendra2 signal. **b**, Representative images of ASPA and SERPINA3N immunostaining. Quantification of ASPA and SERPINA3N-dual labeled cells (arrowheads) and their proportion of total oligodendrocytes (*n* = 4 control and 5 cKO mice, unpaired two-tailed *t* test with Welch’s correction for unequal variance). **c**, Representative image of ASPA and SERPINA3N immunostaining in the cKO tissue, along with quantification of dual-labeled ASPA and SERPINA3N population grouped by mito-Dendra2 presence (normal, low, or absent, *n* = 3 control and 4 cKO mice, unpaired two-tailed *t* test; Welch’s correction for unequal variance was applied to the second and third graphs, left to right). Data are shown as mean ± SEM.

Next, we looked at another stress marker, SERPINA3N, a serine protease inhibitor and a common marker in brain pathologies, implicated, among other things, in cell death regulation and inflammation^80^. More specifically, SERPINA3N is associated with oligodendrocytes upon demyelination^81,82^, and has been shown to mark disease-associated oligodendrocytes in addition to other reactive cells in neurodegenerative conditions and aging^80,83–85^. The number of ASPA-labeled oligodendrocytes positive for SERPINA3N was significantly higher in the cKO tissue, making up approximately 25% of all oligodendrocytes (Fig. 7b). Of note, the majority of these cells (57%) exhibited low levels of mitochondria, whereas the density of oligodendrocytes showing SERPINA3N immunoreactivity and maintaining normal levels of mitochondria was not different from controls (Fig. 7c). The higher likelihood of SERPINA3N presence in oligodendrocytes in cKO tissue that maintained a few mitochondrial puncta as opposed to those displaying complete mitochondrial loss, could suggest a transient increase of SERPINA3N in these cells as a response to stress and survival attempt. Thus, upon FIS1 loss, oligodendrocytes show elevated markers of stress that are also associated with demyelinating conditions, further suggesting impaired homeostasis of these cells and the start of their degeneration.

## DISCUSSION

The concept of prolonged, so-called “dying-back”, death of oligodendrocytes was first proposed decades ago, based on observations from demyelination models and human diseases^15,86,87^. Yet the distinct intracellular processes that underlie this form of degeneration are not fully understood. Here, implementing a precise, optically based approach to induce DNA damage and single-cell demyelination, we reveal that dying oligodendrocytes lose mitochondria within days and persist for weeks to months in their absence. This differs from the other cells in the lineage, which undergo a more stereotyped death progression that involves mitochondrial fragmentation, formation of apoptotic bodies, and cell clearance within hours to days. In addition, we reveal that disruption of a mitochondrial-related gene, *Fis1*, specifically in the oligodendrocyte lineage, leads to oligodendrocyte loss and demyelination but does not impact oligodendrocyte formation. Oligodendrocytes lacking *Fis1* lose mitochondria and display signs of proteotoxic, oxidative, and inflammatory stress, including increased levels of P62 and expression of the disease-associated oligodendrocyte marker, SERPINA3N. Thus, stressed oligodendrocytes, induced either by DNA damage or genetic disruption of a mitochondrial quality control regulator, are characterized by an early loss of mitochondria and persistence of cell soma over prolonged periods. Overall, these results suggest that unique survival and cell death mechanisms exist for oligodendrocytes and that FIS1 and mitochondrial quality control mechanisms are important for maintaining oligodendrocyte homeostasis.

Although 2Phatal induced a similar level of insult across oligodendrocyte lineage cells, it led to different death kinetics and intracellular responses, which point to different modes of death in these cells, as previously suggested^62^. The early stages of the oligodendrocyte lineage are likely to have undergone an apoptotic form of death, supported by observations of mitochondrial fragmentation, cell blebbing, and a rapid death (Fig. 1 and fig. S1). In contrast, persistence of oligodendrocytes’ soma after the drastic collapse of their mitochondria, in addition to the lack of early cellular fragmentation, points to a non-apoptotic death in these cells. In support of this, oligodendrocytes with reduced mitochondria levels in the *Fis1* cKO tissue did not stain positive for CC3, a protease commonly involved in apoptosis^88^, despite showing signs of nuclear changes and cell impairment (fig. S7b). Few or no CC3-positive oligodendrocytes and apoptotic markers have also been seen in other models of demyelination^62^, multiple sclerosis lesions^89,90^, and dying human oligodendrocytes in vitro^39^. Additionally, 2Phatal-targeted oligodendrocytes are likely to maintain their cytoplasmic and membrane integrity, as suggested by retention of tdTomato and, in different scenarios, of a membrane-tethered signal^63^, which also argues against lytic and necrotic cell death in these conditions. Parthanatos is another mode of death associated with oligodendrocytes in some demyelinating contexts^62,75^; however, we did not detect any clear evidence of this pathway in our *Fis1* cKO tissue, despite cells showing hallmarks of death (fig. S7, d and e), suggesting that its activation may be time and context-dependent and leaves open the possibility of alternative mechanisms.

Another form of caspase-independent cell death, where mitochondria are eliminated before cell loss, has been reported in eukaryotic cells in vitro, demonstrating implications of autophagy^91,92^. Similarly, autophagy is likely to play an important role in dying oligodendrocytes seen here, mediating the early degradation of their mitochondria. In support of this, autophagosomes accumulate in metabolically stressed oligodendrocytes^35,93^ and autophagolysosomes are seen engulfing oligodendrocyte mitochondria in myelin mutant animals^94^. Additionally, specific deletion of a critical autophagy gene in mature oligodendrocytes leads to myelin deficits despite preserving oligodendrocyte number^95^, arguing for a role of autophagy in oligodendrocyte and myelin homeostasis. In other contexts, stimulation of autophagy does not rescue the death of stressed oligodendrocytes but exacerbates it instead^96^, suggesting that autophagic mechanisms may act beyond cell maintenance and intersect with cell death. However, the precise relationship between autophagy, survival, and death pathways^97^ in oligodendrocytes is likely context-dependent and would require further investigations.

FIS1 has also been implicated in autophagy and mitochondrial quality control^46,47,98^. Loss of FIS1 impairs mitochondrial respiration, which may induce mitochondrial stress and predispose them to clearance^46^. Mitochondrial clearance upon FIS1 loss is mediated by syntaxin 17, which relocates to mitochondria and recruits the core autophagic machinery^46^. This promotes degradation and eventual loss of mitochondria in cultured cells^46^, consistent with the mitochondrial loss we observe upon depletion of FIS1 in mature oligodendrocytes. In addition, FIS1-depleted mature oligodendrocytes also had elevated P62 levels, likely a sign of overloaded autophagic machinery and impaired autophagy^78^, consistent with other scenarios of FIS1 loss^54^. However, this was not true for all the cells in the cKO tissue, arguing for the transiency of this event in stressed oligodendrocytes that are on their way to death. Additionally, the increase in SERPINA3N presence may have also arisen as a response to oxidative stress^99,100^ in these cells. Its higher occurrence primarily in oligodendrocytes with low mitochondrial levels also suggests a transient expression of this marker, potentially in cells that have entered senescence^99,101^ as a survival attempt.

Depending on the context, FIS1 activity can either be protective against damage and stress, with its loss leading to increased cell death^50,54^ or it can play a pro-apoptotic role instead^58,59^. Based on our results, the former is likely to apply to oligodendrocytes as demonstrated by the eventual loss of FIS1-negative mature oligodendrocytes, going from making up 58% of the oligodendrocyte population 1-month after inducible deletion of *Fis1* in their precursors, to 20% at 3-months, to 1% at 1-year. The decline in overall oligodendrocyte density, however, was not evident at 1 month but became evident at 3 months. Similarly, the oligodendrocyte density did not change at 2 months after inducible deletion of *Fis1* in mature oligodendrocytes despite them showing signs of degeneration, potentially owing to 1) the prolonged death of these cells, which, as seen post 2Phatal, can last up to 10 weeks, and 2) the efficient compensatory mechanism of synchronous remyelination^63^. In support of this, another genetic model targeting a mitochondrial protease in mature oligodendrocytes showed late-stage myelination defects despite early mitochondrial changes^37^, consistent with the observations here. In addition, it is important to note that in our knockout tissue, a mixed population of oligodendrocytes was analyzed, comprising cells at different stages of maturity and non-uniform recombination. This was especially obvious in the *Plp1*-CreER tissue, where tamoxifen injections were spread across 2 months, resulting in mature oligodendrocytes that had undergone recombination between approximately 2 months and 2 weeks before the analysis. This heterogeneity in the deletion timing, coupled with the slow process of mitochondrial turnover and oligodendrocyte degeneration, likely accounts for the variability in mitochondrial content and other morphological and subcellular features observed in these cells, which may correspond to different stages of oligodendrocyte death.

While the occurrence of FIS1-null oligodendrocytes with disrupted mitochondria was observed early on (Fig. 4g) and across all the time points (Fig. 5b and Fig. 6), FIS1-null OPCs maintained their mitochondrial content, at least ∼1-month after Cre recombination. In addition, mitochondrial morphometrics also remained unchanged compared to those in OPCs that escaped the deletion, aligning with the relatively minor role of FIS1 in regulating mitochondrial shape in mammalian cells^48,102^. This is, however, context-dependent^52^, and we cannot exclude the possibility of a mitochondrial remodeling phenotype occurring later, especially given the slow turnover rate of mitochondrial components^103^. Additionally, the depletion of FIS1-null OPCs at the 3-month time point would support a potential transient effect of FIS1 loss in these cells, leading to a change in fate, but given their proliferative nature and high efficiency in restoring their population^70–72^, it did not become evident in our analysis. Overall, this highlights the distinct sensitivities to FIS1 loss in OPCs and oligodendrocytes, which may relate to the different basal levels of FIS1 in these cells as well as their different mitochondrial and population dynamics.

It is important to note that FIS1 has been implicated in other non-mitochondrial functions not explored here, including peroxisomal fission, although its precise involvement remains unclear^52^. Nevertheless, given the pronounced effects induced by FIS1 loss in oligodendrocyte mitochondria, we hypothesize that the observed effects are primarily mediated by altered mitochondrial homeostasis. This, in turn, may have led to elevated reactive oxygen species levels, which cause oxidative damage to cellular proteins, lipids, and DNA^104^. Hence, it is possible that the downstream effects here may converge with what we see by inducing oxidative and DNA damage using 2Phatal (Fig. 2), as supported by observing a similar mitochondrial loss phenotype. In addition, it is also possible that the oxidative stress upon FIS1 loss leads to mitochondrial DNA damage, a common hallmark in demyelinating pathologies^22^, especially given FIS1’s role in preserving mitochondrial DNA integrity^55^. In turn, impairing mitochondrial DNA in mature oligodendrocytes can lead to oligodendrocyte death and demyelination, as shown elsewhere^34^.

Mitochondria have been thought to play less of a prominent role for oligodendrocytes, particularly given that inhibiting oxidative phosphorylation in mature oligodendrocytes does not lead to demyelination^2^. However, the results here suggest that oligodendrocyte health is impaired upon disrupted mitochondrial maintenance despite their prolonged survival. These data suggest distinct mechanisms potentially related to mitochondrial quality control rather than energy production alone. This is consistent with previous studies where genetic disruption of mitochondrial integrity in oligodendrocytes caused oligodendrocyte death and progressive demyelination^34,37^. In addition, hyperactivation of a mitochondrial fission protein in oligodendrocytes induces glycolytic stress, disrupting oligodendrocyte function in Alzheimer’s disease^105^, suggesting that oligodendrocyte mitochondrial impairment may precede pathologies. This might be true among other models of demyelination, in which oligodendrocyte health may be compromised long before myelin loss, glial reactivity, and death markers become apparent^15^ (also supported by findings in Fig. 6g and fig. S7), given that oligodendrocytes’ cell bodies can persist over a month after losing their mitochondria. However, understanding whether an early mitochondrial loss in oligodendrocytes is a common marker of pathologies, would require further longitudinal and cell-level studies in different contexts. Importantly, COX-I, a critical component of the electron transport chain encoded by mitochondrial DNA, was found lacking in oligodendrocytes from multiple sclerosis lesions^23^, and together with the aforementioned evidence, suggest that oligodendrocyte mitochondrial impairment are potentially not an isolated effect of induced DNA damage and *Fis1* deletion but rather a common hallmark of oligodendrocyte degeneration. In support of this, 2Phatal reproduces features of oligodendrocyte degeneration seen spontaneously in aging or in the cuprizone model of demyelination^62,63^, suggesting that the same may apply to the intracellular and mitochondrial dynamics. However, despite cuprizone and experimental autoimmune encephalomyelitis demyelination models showing mitochondrial alterations^30,106^, depletion of mitochondria in these models has not been reported.

Whether persisting oligodendrocytes with mitochondrial deficits remain functional during their prolonged death process, and what supports their maintenance over extended periods, remains unsolved. However, they are likely to lower their metabolism and/or rely on glycolysis^2,35^or on metabolic support from surrounding glia^107–109^, if glycolysis is also compromised following mitochondrial disruptions^105^. Nevertheless, it is likely that mitochondrial impairment in these cells marks the onset of their demise and compromises their potential to support the underlying axons; therefore, the prolonged persistence of these cells could exacerbate axonal vulnerability and pathological burden^110,111^. In light of this, further strategies should consider restoring mitochondrial health in burdened oligodendrocytes to potentially support their function and prevent disease progression.

## METHODS

### Animals

All animal procedures were approved by the Institutional Animal Care and Use Committee at Dartmouth College (protocol #00002158) and were conducted in accordance with the National Institutes of Health Guide for the Care and Use of Laboratory Animals. Animals were housed in a temperature- and humidity-controlled vivarium under a 12 h light/dark cycle and were given ad libitum access to food and water. Both female and male mice were used for all the experiments.

The mouse strains bred in this study include: *Cspg4*-CreER^112^ (JAX #008538); *Plp1*-CreER^113^ (JAX #005975); PhAM (floxed)^114^ (JAX #018385), and Ai9^115^ (JAX #007909), purchased from The Jackson Laboratory, and *Fis1*^loxP^ strain^54^, generously provided by Dr. David Chan’s lab at Caltech. PhAM mice express a mitochondrial matrix-targeted Dendra2 fluorescent protein, and Ai9 mice express tdTomato fluorescent protein following Cre-mediated recombination. *Cspg4*-CreER mice label OPCs and mural cells upon tamoxifen-induced Cre-mediated recombination. Since this study was focused on OPCs and their progeny, labeled mural cells were identified based on their location and morphology and excluded from any analysis. *Plp1*-CreER mice label myelinating cells upon tamoxifen-induced Cre-mediated recombination. *Fis1*^loxP^ mice undergo Cre-mediated excision of the floxed exon 2 of the *Fis1* gene, which generates a null allele due to a frameshift mutation^54^. Genotyping of the *Fis1*^loxP^ allele was done via Transnetyx. Intraperitoneal injections of tamoxifen (Sigma-Aldrich #T2859) dissolved in corn oil were performed to induce Cre-mediated recombination. Intraperitoneal injections of EdU (50 mg/kg, Vector Labs #1149) prepared in sterile PBS were performed to detect proliferating cells. Three EdU injections were administered at 4-hour intervals, and mice were perfused 24 hours following the first injection.

For 2Phatal experiments, triple transgenic *Cspg4*-CreER; Ai9/PhAM mice were used. 2Phatal of newly formed oligodendrocytes was performed in young adult mice (∼2 months-old) to increase the likelihood of capturing recombined cells at an early stage of maturation. 2Phatal of mature oligodendrocytes was done in older adult mice (∼7 months old and ∼10 months old for the multifocal 2Phatal). 2Phatal of OPCs was done across all ages. A single dose of tamoxifen (0.5 mg) was injected at P18.

For conditional deletion of *Fis1* in OPCs and their progeny, *Cspg4*-CreER; *Fis1*^loxP/loxP^ mice with or without mitochondrial and cytoplasmic reporters were used, depending on availability. Littermates with matching genotype and wild-type *Fis1* were used as controls. For experiments involving one-year-old animals, *Fis1* heterozygous mice were used as controls due to the limited availability of littermates carrying the wild-type *Fis1* allele. Two doses of tamoxifen (0.1 mg) were injected at P5 and P7 for both floxed and control mice. Tissue collection was done approximately at 1 month (P29-P37), 3 months (P100-102), and 12 months (P365) after tamoxifen injection.

For conditional deletion of *Fis1* in myelinating oligodendrocytes, *Plp1*-CreER; *Fis1*^loxP/loxP^ with or without mitochondrial reporter were used, depending on availability. Littermates with matching genotype and wild-type *Fis1* were used as controls. Mice with mitochondrial reporter were used for analysis that involved mitochondrial quantification, whereas all available relevant genotypes were used for cell density measurements and oligodendrocyte lineage progression analysis. Eight doses of tamoxifen (1 mg) were injected over a period of 6 weeks, starting at P65-P70, for both floxed and control mice. Tamoxifen was given every 2-3 days across 3 cycles separated by 2-week intervals. Tissue collection was done ∼2-3 weeks after the last injection, corresponding to 2 months after the first injection (P125-P130).

### Surgical Procedures

To create an optically accessible region for intravital imaging, cranial windows were implanted over the somatosensory cortex. Mice were first anesthetized via an intraperitoneal injection of ketamine (100 mg/kg) and xylazine (10 mg/kg) solution. Toe-pinch reflex testing was performed to ensure adequate anesthesia, and a single dose of carprofen analgesic (5 mg/kg) was subcutaneously administered before the procedure. An ophthalmic ointment was applied to the eyes to prevent them from drying. The surgical area was shaved, sterilized, and the underlying skull was exposed. A custom metal nut was affixed to the rostral portion of the skull to allow stable head fixation during surgery and imaging. Using a high-speed drill, a 3×3 mm craniotomy was performed over the somatosensory cortex in both hemispheres, and the underlying dura was removed. Hoechst 33342, a DNA-binding fluorescent dye (ThermoFisher #H3570; 0.05 mg/ml in PBS) was topically applied to the exposed pial surface to label cortical nuclei for 2Phatal experiments. A #0 cover glass was placed over the craniotomy, and the surrounding skull was sealed with dental cement. A second dose of carprofen analgesic was injected at the end of the surgical procedure, as well as 24 and 48 hours after. Mice were monitored throughout the procedure and returned to the recovery cage.

### Imaging

For intravital imaging, mice were anesthetized with isoflurane (2% for induction and 0.8-1.2% during imaging), head-fixed, and placed under the microscope. Single time point images were taken the day of the surgery using an upright laser-scanning confocal (Leica SP8 with LasX software version 3.5.7) with a 20x water-immersion objective (Leica, NA 1.0). The tdTomato and mito-Dendra2 were excited respectively by 488 and 552 nm laser wavelengths. 2Phatal experiments were performed the day after the surgery to allow for optimal nuclear labeling, using an upright laser-scanning 2-photon microscope (Bruker Ultima with Prairie Software version 5.4) with an InSight X3 femtosecond pulsed laser (Spectra Physics) and a 20x water-immersion objective (Zeiss, NA 1.0). The tdTomato, mito-Dendra2, and Hoechst 33342 were excited respectively by 1040, 920, and 775 nm laser wavelengths. Z-stacks were acquired using interlaced scanning over 50-100 µm depth in the somatosensory cortex. Longitudinal imaging was done at 3, 5, and 7 days post 2Phatal and continued weekly for 2-3 months until the cells were cleared. Stable landmarks such as vasculature and/or myelin were used as a reference to relocate the same field of view.

Fixed brain slices were imaged over the cortex between the pial surface and deeper cortical regions above the white matter using a Leica SP8 laser-scanning confocal with 20x air (Leica NA, 0.75), 20x water-immersion (Leica, NA 1.0), or 63x oil-immersion (Leica, NA 1.4) objectives or with a Nikon SoRa spinning disk confocal (with NIS-Elements AR software version 6.10.02) with 20x air (Nikon, NA 0.8) or 40x oil-immersion (Nikon, NA 1.3) objectives. The following excitation wavelengths were used: 405, 488, 552, and 638 nm. Sequential imaging was done from longest to shortest wavelength. Images were taken across 2-4 coronal sections per animal, with 1-3 fields of view per hemisphere in each section. The image settings and regions imaged were kept consistent for control and experimental conditions. To prevent bias, channels for mitochondrial (mito-Dendra2, TOM20) or other subcellular labels (P62) were off during selection of the fields of view.

### Single-Cell and Multifocal 2Phatal

2Phatal was performed to induce targeted death of single cells in the oligodendrocyte lineage following DNA damage. First, the nuclear labeling dye Hoechst 33342 was applied during the cranial window preparation as described above. After allowing 24 hours for recovery after surgery and uniform nuclear labeling, sequential images of tdTomato, mito-Dendra2, and Hoechst fluorescence were taken with the 2Photon microscope to visualize the cells and their corresponding nuclei. A random subset of cells that had mitochondrial labeling was selected at each stage in each field of view. To identify the cellular maturation stage, unique mitochondrial and morphological features were used. OPCs were identified based on their branched, complex processes and a heterogeneous distribution of mitochondria in the cell. Newly formed oligodendrocytes were identified based on the presence of newly formed sheaths, their large soma size, high mitochondrial content in the periphery, and either high or low mitochondrial content in the soma, which indicates how recently the cell differentiated^60^. Oligodendrocytes were identified based on a smaller soma size with high mitochondrial content, thin processes, and low mitochondrial presence in the periphery. Once targeted cells were selected and identified, an ROI of 10×10 pixels was drawn over each corresponding nuclei and away from nearby mitochondrial signal. To induce photobleaching of the nuclear dye, the laser wavelength was set at 775 nm, the dwell time at 100 µs, and a time series of 150 scans was taken over the defined region, lasting a total of 1.2 seconds. Untargeted, neighboring cells were used as control. After 2Phatal, the field of view was re-imaged to ensure that no visible damage occurred to the cell cytoplasm or its mitochondria. Longitudinal imaging was done periodically within the first week of 2Phatal, followed by weekly imaging until the targeted cells disappeared. This was evidenced by the loss of tdTomato signal and classified as the day of cell death.

Multifocal 2Phatal was performed to target a larger number of mature oligodendrocytes using the method described above. Approximately 400-500 mature oligodendrocytes, which started with normal mito-Dendra2 labeling, were targeted per mouse across 10-12 fields of view per hemisphere. Follow-up imaging was performed within the first week of 2Phatal and the day before tissue collection to confirm successful targeting. Tissue collection was performed at 28 days post-2Phatal. The tissue was processed, and coronal sections were stained for the mitochondrial marker TOM20, oligodendrocyte marker CNP, and DNA marker Hoechst 33342. Fixed tissue imaging was done over the somatosensory cortex corresponding to the site of in vivo 2Phatal. Targeted cells were identified based on their nuclear appearance: small, and with indiscernible nucleoli, while being blinded to endogenous Dendra2 labeling and TOM20 immunolabeling of mitochondria. Control cells were identified based on their unperturbed nuclear morphology.

### Intravital Imaging Analysis

After 2Phatal experiments, analysis was performed on a randomly selected set of cells that were successfully targeted and underwent death, along with control cells from the same environment. To formally confirm the stage of the targeted cells, area and mito-Dendra2 intensity were measured within a circular or elliptical ROI drawn around their somas. All analysis was performed on single optical sections at the middle of the cell volume. Only the tdTomato was used as a reference for the soma outline while being blinded to the mito-Dendra2 channel. To detect early changes in mitochondrial shape, paired analysis was performed at baseline (before 2Phatal) and 3 days after 2Phatal. First, the mito-Dendra2 signal within the soma ROIs was thresholded, and mitochondrial ROIs were automatically generated using the “Analyze Particles” command in FIJI. Measurement of circularity (located in FIJI’s shape descriptors) was done for each mitochondrial ROI and averaged per cell. To detect changes in tdTomato and mito-Dendra2 fluorescence over time, analysis was done at baseline, immediately after 2Phatal, at 3, 5, and 7 days, and then weekly following 2Phatal until the time point before the tdTomato signal was lost, which for some cells occurred up to 70 days post-targeting. Mean intensities of tdTomato and mito-Dendra2 were measured within the soma ROIs. ROI sizes were consistent across all time points to ensure that intensity measurements were comparable throughout. A ratio of mito-Dendra2 to tdTomato mean fluorescence intensity was taken within each ROI over time and normalized to the baseline time point to quantify relative fluorescence changes over time.

### Tissue Processing and Immunohistochemistry

The brain tissue was collected and processed at the end of each experiment. First, the animals were fully anesthetized via ketamine (100 mg/kg) and xylazine (10 mg/kg) injections and transcardially perfused with 4% paraformaldehyde solution in 1x PBS. The brains were then dissected and stored in 4% paraformaldehyde solution at 4°C for the first 24 hours before being transferred to 1x PBS. Next, brains were sectioned coronally at 75 µm, and the generated free-floating sections were placed in glycerol-based cryogenic solution for long-term storage at −20°C.

For immunostaining, first, the sections were washed three times in 1x PBS for 10 minutes each. Antigen retrieval was performed by placing the sections for 2 minutes in a 95°C pre-heated buffer solution consisting of 10 mM Tris, 1 mM EDTA, and 0.05% Tween 20 in 1x PBS at pH 9. The sections were brought to room temperature in the same solution before being transferred to primary antibody diluted in blocking and permeabilization solution (0.3% Triton X-100 and 1% Bovine Serum Albumin in 1x PBS) and placed on a rotating shaker at room temperature for 24 hours. The primary antibodies used include: guinea pig anti-CNP (1:500, Synaptic Systems, Catalog #: 355004), mouse anti-CAII (1:500, Santa Cruz Biotechnology, #sc-48351), rabbit anti-Aspartoacylase (1:1000, GeneTex, #GTX113389), rat anti-MBP (1:100, BioRad, #aa82-87), rabbit anti-BCAS1 (1:750, Synaptic Systems, #445 003), goat anti-PDGFRA (1:1000, R&D Systems, #AF1062), rabbit anti-IBA1 (1:500, FUJIFILM Wako, #019-19741), rabbit anti-GFAP (1:500, Sigma-Aldrich, #G9269), goat anti-human/mouse caspase-3 (1:200, R&D Systems, #AF-605-SP), rabbit anti-cleaved caspase 3 (1:700, Cell Signaling Technology, #9661S), goat anti-SERPINA3N (1:300, R&D Systems, AF4709), rabbit anti-KI67 (1:250, Abcam, #Ab16667), rabbit anti-TOM20 (1:200, Abcam, #ab78547), rabbit anti-FIS1 (1:200, Proteintech, #10956-1-AP), rabbit anti-P62 (1:500, Abcam, #ab91526). Following primary antibody incubation, the sections were washed three times in PBS and incubated for 2 hours at room temperature in secondary antibody diluted in blocking and permeabilization solution. Secondary antibodies were conjugated as needed to Alexa Fluor 488, 555, or 647, or to CF555 or CF647 (1:500; Thermo Fisher Scientific, Jackson ImmunoResearch, or Biotium). After two more washes in PBS, the sections were incubated for 20 minutes at room temperature in a Hoechst 33342 solution (10 mg/ml stock) diluted at 1:2000 in 0.3% Triton X-100 in 1x PBS. After a final PBS wash, the sections were then mounted with ProLong Diamond Antifade Mountant (Thermo Fisher Scientific, #P36970) on a glass slide and covered with a #1.5 coverslip.

The same steps were followed for PAR labeling, except that the slices were incubated with the primary antibody solution containing the anti–PAR binding reagent (1:500, Sigma-Aldrich #MABE1031) overnight at 4°C instead of at room temperature. The rabbit Fc tag on the PAR reagent was detected with an anti-rabbit secondary antibody. If needed to quench the endogenous fluorophores, sections were incubated for 1 hour at room temperature in solution containing 10 mM copper (II) sulfate and 50 mM ammonium acetate solution at pH 5. To quantify cell proliferation, immunohistochemistry was followed by EdU detection using the Click-iT kit (ThermoFisher Scientific, #C10340) and Alexa Fluor 647 dye following the manufacturer’s instructions.

### Fixed Tissue Analysis

Analyses were performed on 63x, 40x, or 20x images, depending on the resolution required. TOM20 immunostaining analysis was performed on single optical z-sections to avoid any overlap of the broad TOM20 signal across the z-stack. The sections corresponded to the middle of the cell volume and were selected based on CNP immunolabeling. An ROI was drawn over the somas and nuclei of targeted and control cells using the selection brush tool in FIJI. When defining the soma, only the CNP channel was used for reference to avoid any bias from the mitochondrial signal. Note that occasionally TOM20 signals of uncertain cellular origin would be included in the ROIs due to blinding. Measurements of soma and nucleus area, along with the mean intensity of TOM20 and Dendra2 signal, were taken per cell. Both intensities were corrected for background, calculated as the average intensity from 3 fixed-sized ROIs in areas without detectable signal. The same steps were also followed for analysis of P62 mean intensity within cell somas, with analysis done on maximum intensity projections of 3 sections to capture a larger volume.

To analyze FIS1 fluorescence levels across the oligodendrocyte lineage, first, a group of PDGFRA- and CAII-labeled cells was randomly selected while being blinded to the FIS1 immunostaining. Analysis was done on cropped single optical z-sections that had both soma and process mitochondria from each cell type in focus. Measurements were made within automatically generated mitochondrial ROIs to account for mitochondrial content. Individual mitochondrial ROIs per cell were combined into one, and the mean intensity for FIS1 immunostaining within the generated ROI was measured and corrected for background.

*Fis1* cell-specific deletion was confirmed by analyzing the lack of FIS1 immunostaining in these cells using manual and automated approaches. For the first round of analysis (Fig. 4, d and e, and fig. S4a) high-resolution images of PDGFRA- and CAII-labeled cells were randomly selected while being blinded to the FIS1 immunostaining to avoid any bias. Following up, a qualitative assessment was done to confirm the overlap of cell-specific mito-Dendra2 and FIS1 labels. If no overlap was seen, the cell was denoted as FIS1-negative. After identifying the criteria for a positive FIS1 signal from this initial analysis, subsequent analysis (Figs. 5 and 6) was done without reference to the mito-Dendra2 channel to include a larger population of cells regardless of mito-Dendra2 recombination. This qualitative analysis was then validated by an automated process done on low-magnification images. For this, first the maximum intensity projections were prepared and the CAII-channel was thresholded. The thresholded images were loaded in TrackMate^116^ (version 7.7.2), applying a LoG detector with an estimated object diameter of 8-11 µm (depending on image magnification) that corresponded to the CAII-labeled cells soma size and a quality threshold of 0.6-1.2 (optimized depending on image quality). The generated spots depicting CAII-labeled somas were then exported as ROIs. Automatically generated ROIs were overlaid with the corresponding thresholded FIS1 channel, and the area fraction of FIS1 signal within the ROIs was measured. Baseline FIS1 area fraction levels in the control tissue were used to determine a threshold that would distinguish between positive and negative FIS1 signals. Note that occasionally false negative signals would be generated for some cells in the control tissue, potentially due to the FIS1 signal not being included in the maximum projection. After confirming that the results between manual and automatic approaches were comparable and reliable for the data in Figs. 4 and 5, only the automatic approach was used for the results in Fig. 6.

To analyze the effect of FIS1 deletion in mitochondrial and OPC morphology, analysis was done on a randomly selected set of OPCs in the cKO tissue, later categorized into cells with positive or negative FIS1 signal as determined by the FIS1 immunostaining. Cells with a positive FIS1 signal were used as internal controls. Maximum projections of the cells of interest were processed, thresholded, and cleared of other interfering cell structures from the environment. ROIs for the cytoplasm and mitochondria were automatically generated using FIJI’s Analyze Particles command with PDGFRA and mito-Dendra2 channels as reference. Area fraction of mitochondrial signal within the cytoplasm ROI was measured as a proxy for mitochondrial content. Fit ellipse and shape descriptors commands in FIJI were used to measure mitochondria size and shape. The major and minor axes of the fitted ellipse for each mitochondrial ROIs were extracted and used as a proxy for mitochondrial length and width, respectively.

The same images were used to perform SNT “Sholl Analysis” ^117^ and quantify the branching of PDGFRA-positive cell processes. A straight line was drawn from the center of the cell soma to the most distal point of the image to define the center of the analysis. The start radius was set at 3 µm, the step size at 1 µm, and the end radius at 75 µm. The total number of intersections was summed per cell.

Cell density analysis was performed on maximum projections of fixed volumes. ASPA- and CAII-labeled cell density in Figs. 4f, 6g, and fig. S4c was done automatically by counting the ROIs generated using TrackMate as described above. The round soma shape of oligodendrocytes and their uniformity allowed for their reliable automatic detection. The rest of the cell counts were done manually. For cell death counts, only the CC3-labeled cells that also displayed condensed nuclei were counted to avoid counting any cellular debris. In fig. S3d, dying cells were counted in intravital images and identified by fragmented, apoptotic-like, tdTomato signal. To quantify territorial coverage for MBP, GFAP, and IBA1 immunolabeling, maximum projections were prepared, thresholded, and the area fraction was calculated. To quantify the overlap between MBP and BCAS1, first, ROIs were generated from the thresholded MBP channel using “Create selection” command in FIJI. The area fraction of the thresholded BCAS1 signal was measured within the generated ROIs.

Automated and manual approaches were used to determine the levels of mito-Dendra2 per cell. For the automated approach, oligodendrocyte soma ROIs were automatically generated, and the area fraction of thresholded mito-Dendra2 signal was measured. Cells were classified as having normal, low, or no mito-Dendra2 signal when area fraction values within ROIs were respectively above 10%, between 2 and 10%, or below 2%. Manual assessment of mito-Dendra2 levels per cell was done qualitatively by independent experimenters and yielded consistent results with automated analysis.

### Statistical Analysis

All statistical analyses were performed using GraphPad Prism (version 10.4.2) or Excel. Data are displayed as mean ± SEM. For 2Phatal analysis, each data point represents a cell, given the single-cell specificity of the technique. For comparisons between control and cKO groups, each data point represents the average per animal, calculated by averaging measurements obtained per cell or per field of view. Given the cell-to-cell variability in these groups, cell-level data points are also shown as indicated. Sample size determination was based on previous publications^63,110^. Normal distribution was assumed but not formally tested unless otherwise specified. For comparisons between two groups, paired or unpaired two-tailed Student’s *t* test was used. *F* test was run to compare variances, and if variances were significantly different, Welch’s correction was used. For comparisons between multiple groups, the Brown-Forsythe test was run to compare variances, and Welch ANOVA with Dunnett’s T3 multiple comparisons test was used for unequal variances. For datasets with zero within-group variance, Kruskal Wallis with Dunn’s T3 multiple comparisons test was applied. The analyzed cells were randomly selected while being blinded to any subcellular label to avoid any bias. The experimenter was also blinded to the animal genotype. No animals were excluded from the statistical analysis.

## Acknowledgments

We thank Drs: David Chan and Hsiuchen Chen from Caltech for sharing the *Fis1*^loxP^ line with us. We also thank the Hill lab members for helpful discussions on the project.

## Author Contributions

XB and RAH conceived and designed the study and secured funding. XB performed most of the experiments, data analysis, and visualization, and wrote the original manuscript. SZG contributed to staining and imaging for *Fis1*-related experiments, analyzed the 3-month timepoint, and performed KI67^+^ and IBA1^+^ cell counts. YK performed PAR staining, related figure and analysis, and manual quantification of mito-Dendra2 presence in CNP^+^ cells. RAH and XB performed cranial windows. All authors edited and approved the manuscript.

## Data and materials availability

All data are available upon request.

