## Supplementary Figures for "Dying oligodendrocytes persist without mitochondria"

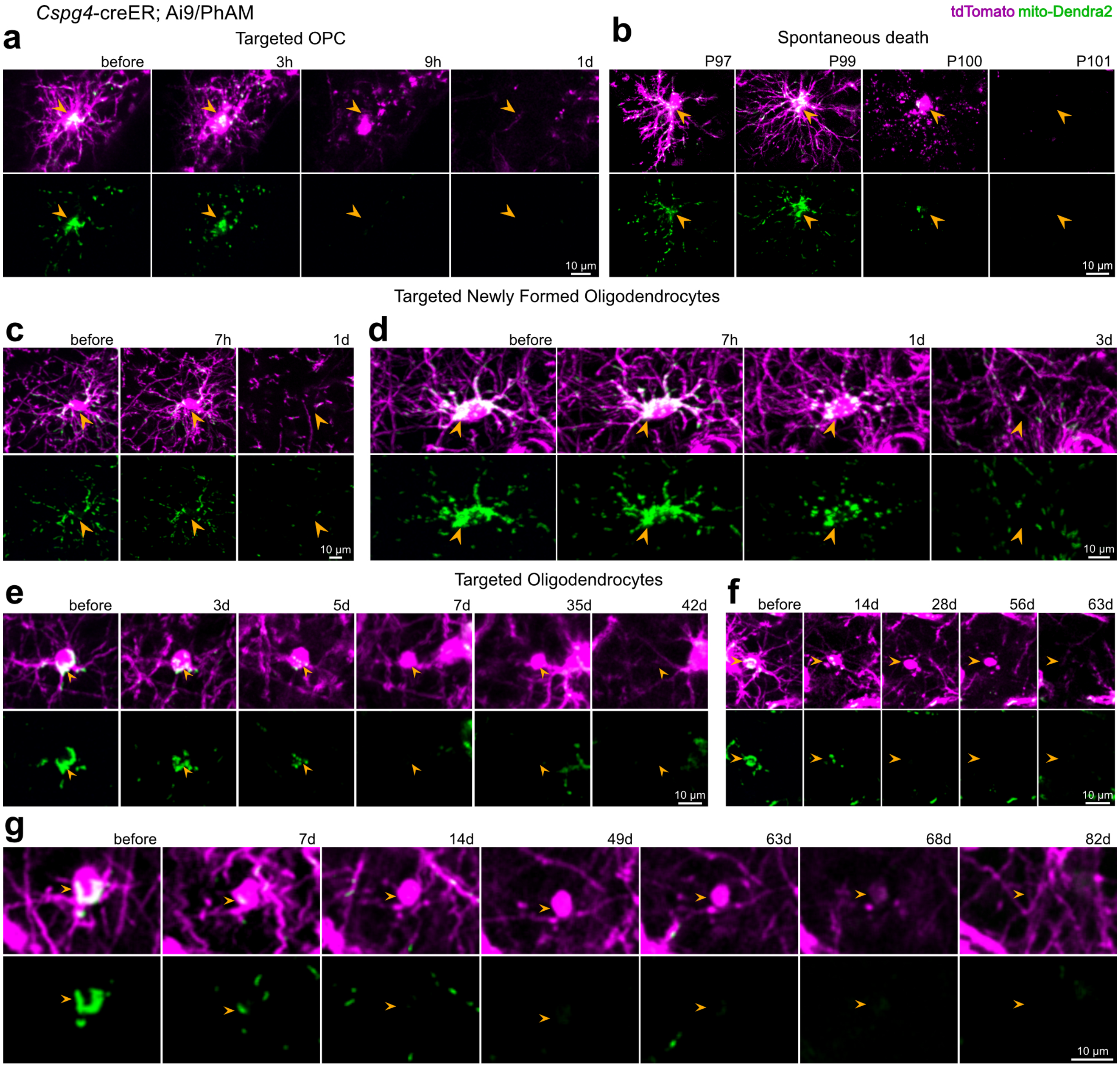


**Figure S1. Cell death and mitochondrial dynamics across the oligodendrocyte lineage.**
**a,** 2Phatal targeted OPC undergoing mitochondrial and cellular fragmentation starting at 3 hours after 2Phatal and getting cleared within a day. **b,** OPC undergoing spontaneous death showing similar features of a 2Phatal targeted OPC. **c, d,** 2Phatal targeted newly formed oligodendrocytes undergoing death following the onset of mitochondrial disruptions, and getting cleared within 1 **(c)** and 3 **(d)** days. **e, f,** 2Phatal targeted mature oligodendrocytes undergoing death and persisting for at least 4 weeks without mitochondria. Cells are cleared out by day 42 **(e)** and 63 **(f)** following 2Phatal. **g,** 2Phatal targeted mature oligodendrocytes undergoing death and persisting for at least 8 weeks without mitochondria. Note restoration of myelin coverage by day 82. Arrowheads point at dying cells.

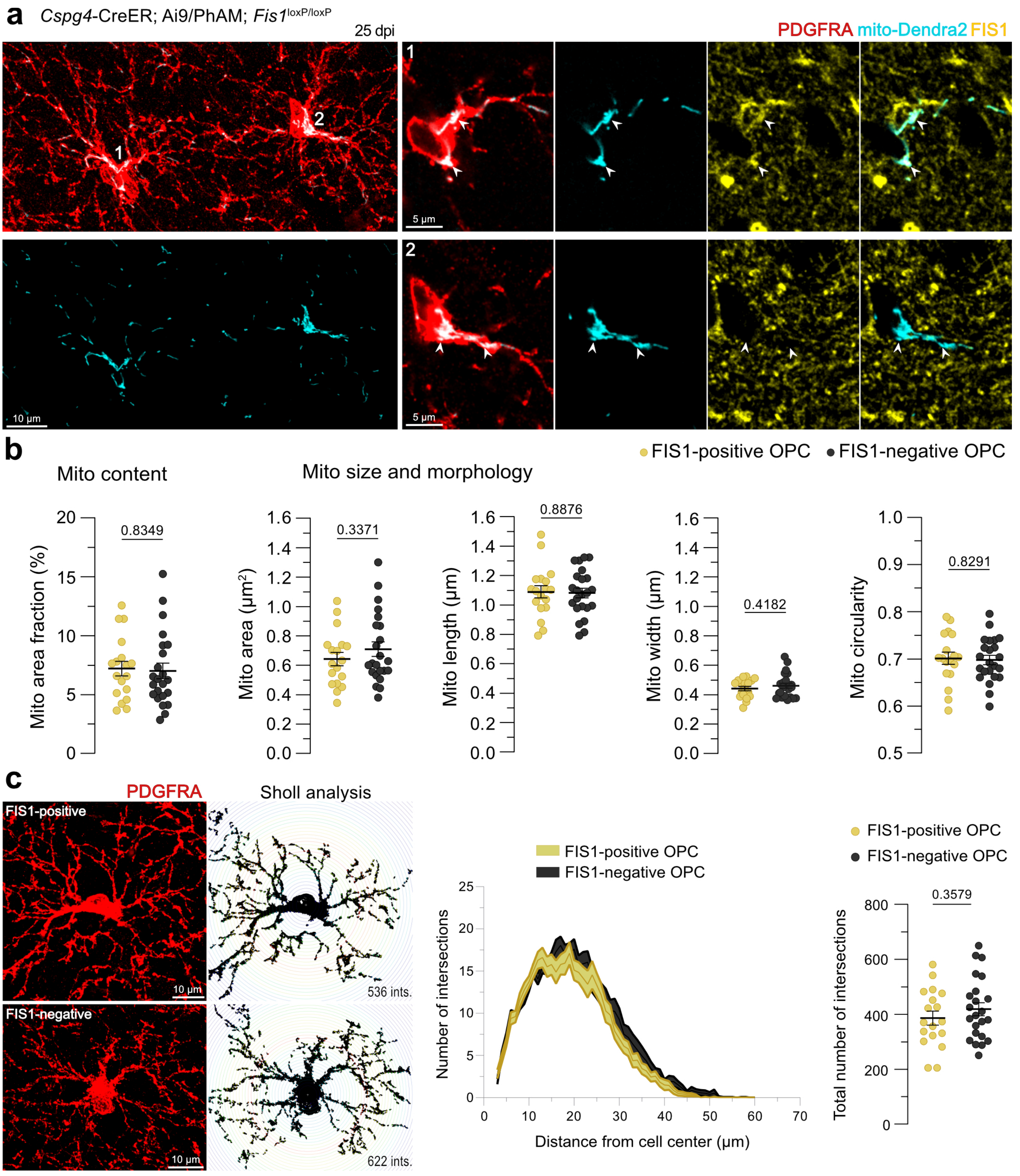


**Figure S2. Mitochondrial morphometrics are not different in FIS1-negative OPCs.**

**a,** A FIS1-positive PDGFRA$^+$ cell (1) serving as an internal control, and a FIS1-negative PDGFRA$^+$ cell (2) in the cKO tissue. Arrowheads point at mitochondria. **b,** Mitochondrial content, size, and morphology quantified as mitochondria to cytoplasm area fraction, mitochondrial area, length, width, and circularity. **c,** Examples of Sholl analysis performed on a FIS1-positive and a FIS1-negative PDGFRA$^+$ cell. The number of intersections is shown as a function of distance from the center of the cell. Quantification of the total number of intersections per cell. In **(b)** and **(c)**, $n$ = 18 FIS1-positive PDGFRA$^+$ cells and 23 FIS1-negative PDGFRA$^+$ cells from 6 cKO mice, unpaired two-tailed $t$ test. Data are shown as mean ± SEM.

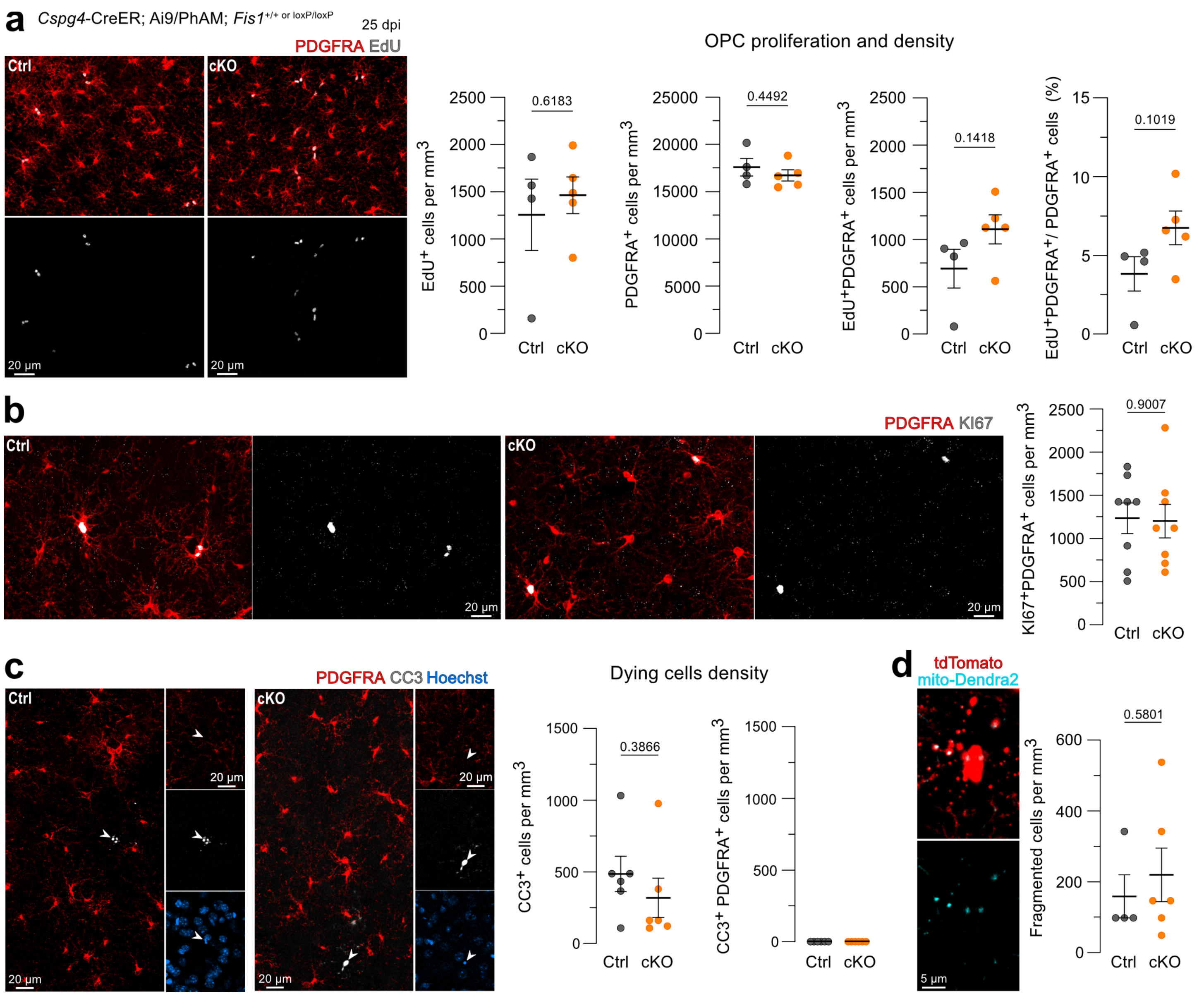


**Figure S3.** ***Fis1*** **loss does not change OPC population homeostasis.**

**a,** Representative fields of view showing PDGFRA staining and EdU labeling. Quantification of the total number of EdU$^+$, PDGFRA$^+$, and dual-labeled EdU$^+$ PDGFRA$^+$ cells along with the percentage of the dual-labeled cells across total PDGFRA$^+$ cells ($n$ = 4 control and 5 cKO mice, unpaired two-tailed $t$ test). **b,** Representative images and quantification of PDGFRA$^+$ cells with KI67 labeling ($n$ = 8 control and 8 cKO mice, unpaired two-tailed $t$ test). **c,** PDGFRA and CC3 staining in the cerebral cortex. Arrowheads point at CC3$^+$ cells with condensed nuclei. Note the lack of PDGFRA and CC3 co-labeling. Quantification of the total number of CC3$^+$ cells and lack of CC3$^+$ PDGFRA$^+$ cells in either condition ($n$ = 6 control and 6 cKO mice, unpaired two-tailed $t$ test). **d,** Representative in vivo image showing fragmented mitochondria and apoptotic bodies in a dying oligodendrocyte lineage cell. Quantification of dying cell density ($n$ = 4 control and 6 cKO mice, unpaired two-tailed $t$ test). Data are shown as mean ± SEM.

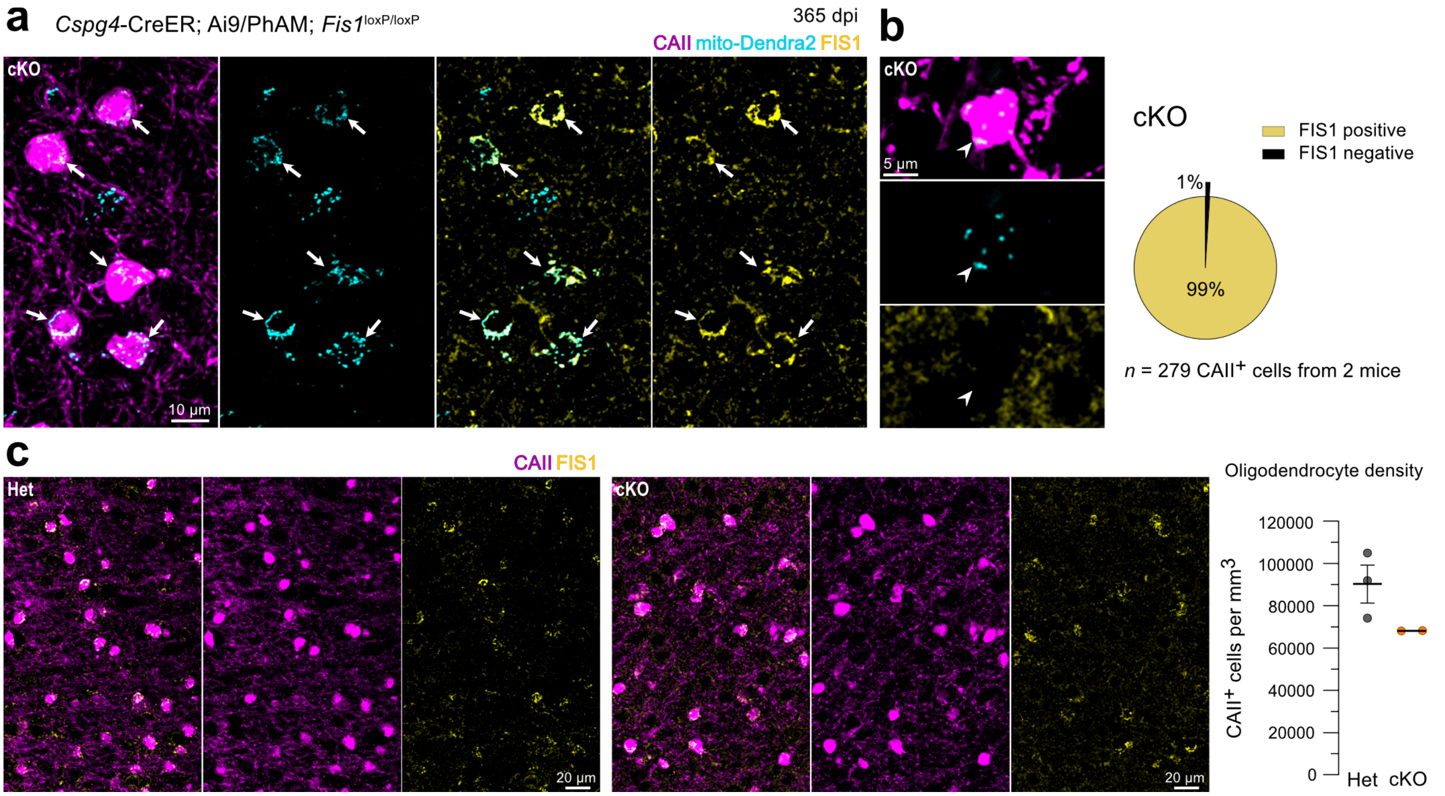


**Figure S4. Progressive loss of *Fis1*-negative oligodendrocytes one year post Cre recombination.**

**a,** Representative images of oligodendrocytes and FIS1 staining 1 year after Cre recombination in the cKO tissue. **b,** Example of a FIS1-negative oligodendrocyte and percentage of this population across all analyzed oligodendrocytes in the tissue. In **(a)** and **(b),** arrows point at FIS1-positive oligodendrocytes, and arrowheads point at FIS1-negative oligodendrocytes. **c,** Representative images and quantification of CAII$^{+}$ cell density (*n* = 3 heterozygous and 2 cKO mice). Data are shown as mean ± SEM.

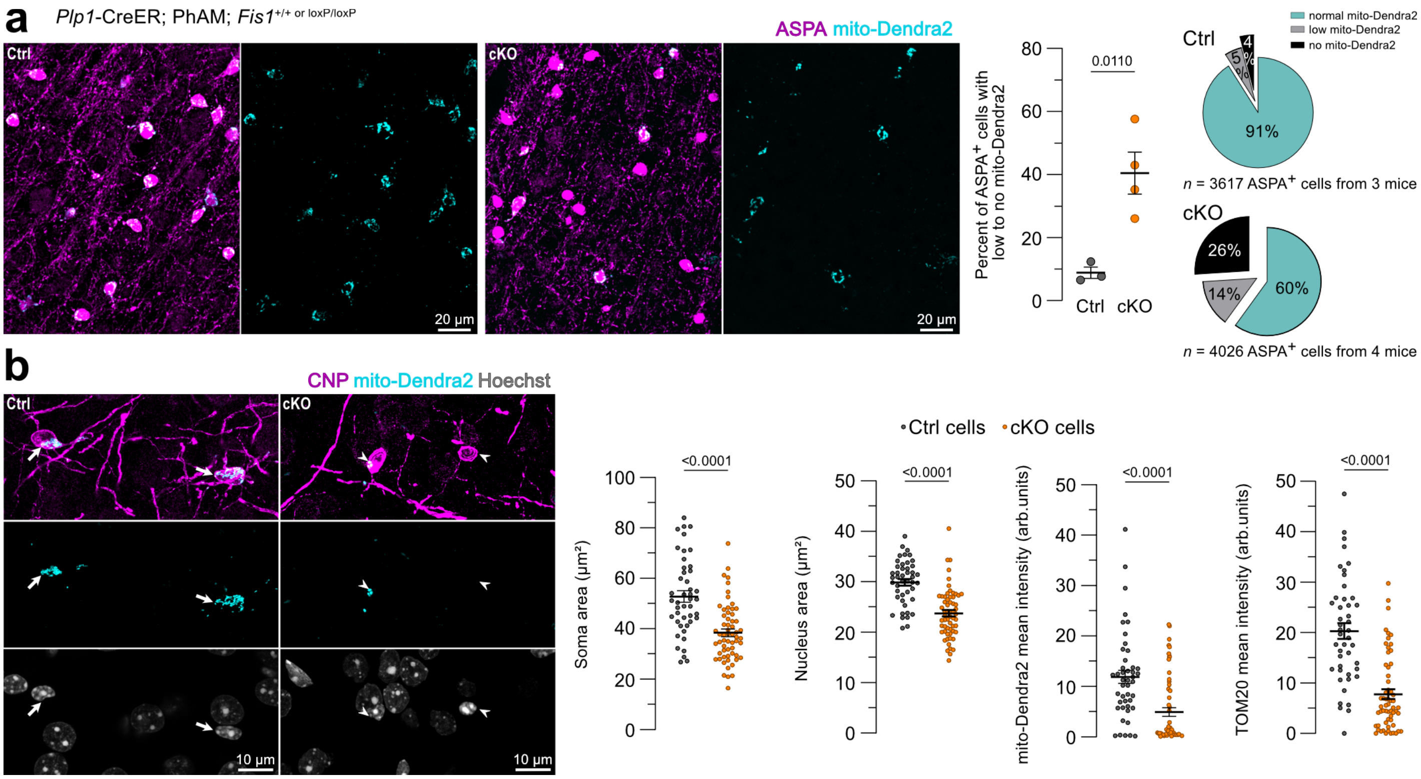


**Figure S5. Mitochondrial loss, morphological changes, and cell heterogeneity after *Fis1* deletion in oligodendrocytes.**
**a,** Representative fields of view showing ASPA-labeled oligodendrocytes and their mitochondria in control and cKO mice. Automated quantification of the percentage of ASPA$^+$ cells with low or no mito-Dendra2 signal ($n$ = 3 control and 4 cKO mice, unpaired two-tailed $t$ test). **b,** Representative images of oligodendrocytes with normal mitochondrial content and normal nuclear morphology in the control (arrows), and oligodendrocytes with low (left) and no (right) mitochondrial content and condensed nuclei in the cKO tissue (arrowheads). Note: the cell lacking mitochondria in the cKO image is the same cell shown in Fig. 7a. Quantification of soma area, nucleus area, and soma mito-Dendra2 and TOM20 mean fluorescence intensity per cell ($n$ = 45 cells from 3 control and 60 cells from 4 cKO mice, unpaired two-tailed $t$ test; Welch's correction for unequal variance was applied to the first, third, and fourth graph, left to right). Data are shown as mean ± SEM.

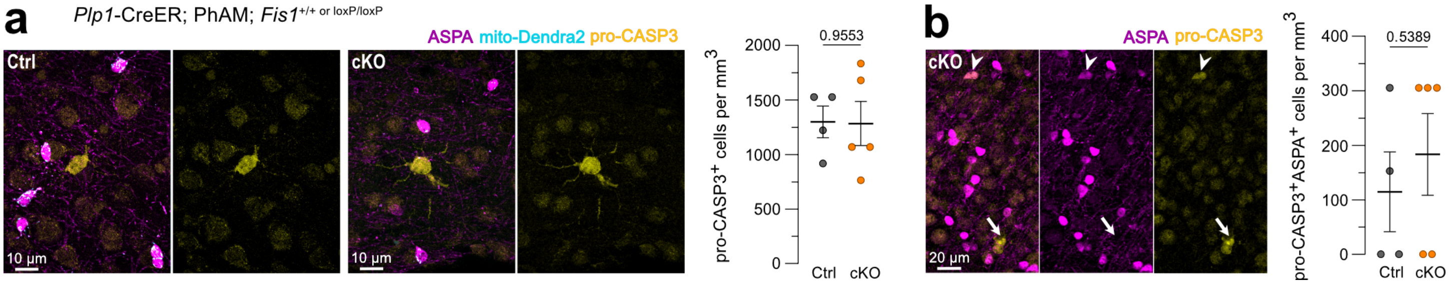


**Figure S6. *Fis1* deletion in oligodendrocytes does not change the density of differentiating oligodendrocytes.**
**a,** Representative images of ASPA and pro-CASP3 staining. Quantification of the total pro-$CASP3^+$ **(a)** and dual-labeled pro-$CASP3^+ASPA^+$ cells **(b),** (*n* = 4 control and 5 cKO mice, unpaired two-tailed *t* test). Arrowhead points at a pro-$CASP3^+ASPA^+$ cell, and the arrow points at a pro-$CASP3^+ASPA^-$ cell. Data are shown as mean ± SEM.

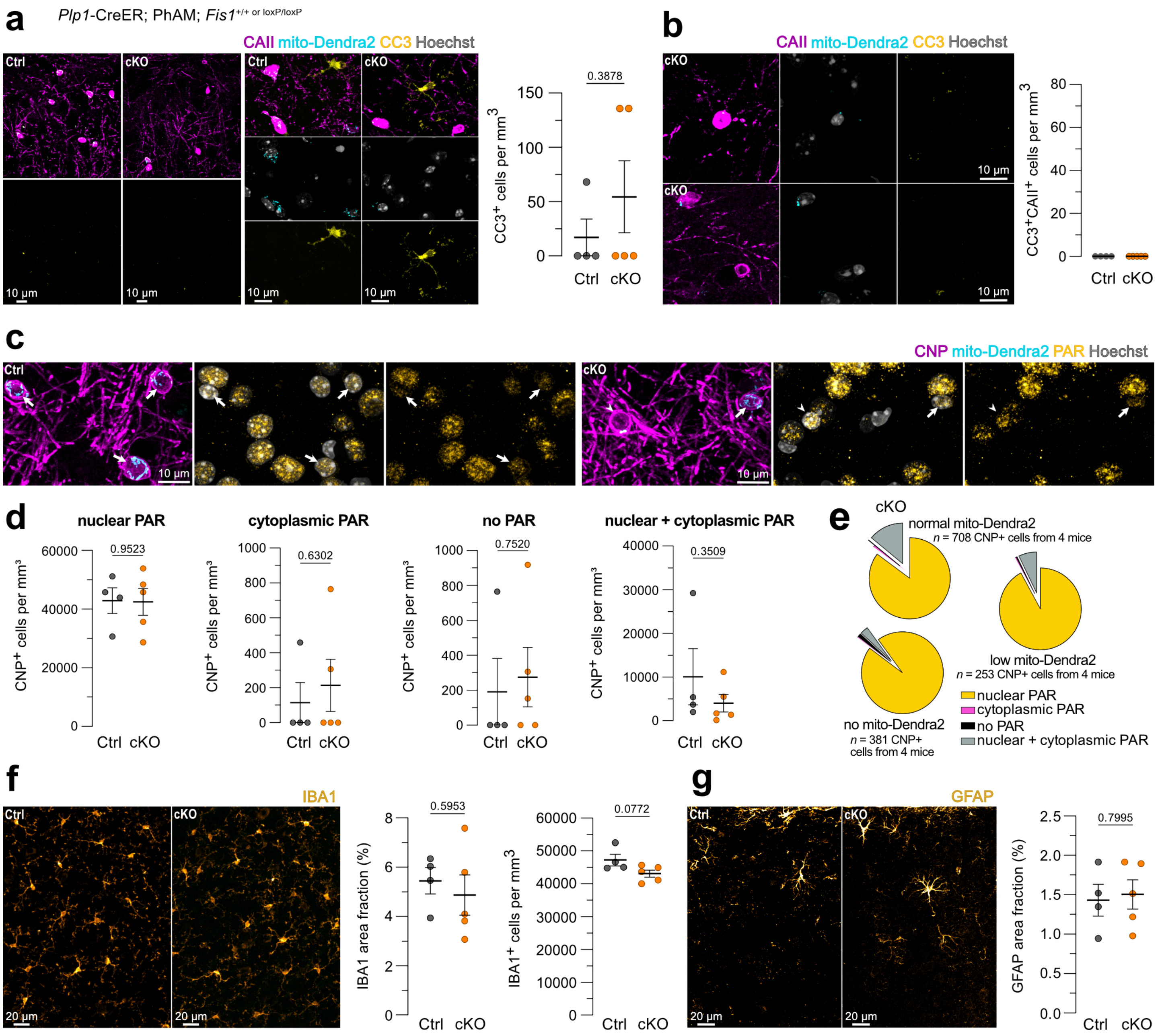


**Figure S7. Lack of apoptotic, parthanatos, and glial reactivity markers after *Fis1* deletion in mature oligodendrocytes.**
**a,** Representative images of CAII and CC3 staining. High magnification images show CC3$^+$CAII$^-$ cells in the territory. Quantification of total CC3$^+$ cells. **b,** Representative images and quantification showing lack of CC3$^+$CAII$^+$ population, despite CAII$^+$ cells displaying abnormal mitochondrial content and nuclear morphology ($n$ = 4 control and 5 cKO mice). **c,** Representative images of PAR staining and localization in the cell. **d,** Quantification of CNP$^+$ cell densities grouped according to PAR localization (nuclear, cytoplasmic, absent, or nuclear and cytoplasmic). **e,** Pie charts showing PAR localization grouped according to the mitochondrial content in the cell (normal, low, absent) in the cKO tissue. **f,** Immunostaining and quantification of IBA1$^+$ area coverage and cell density. **g,** Immunostaining and quantification of GFAP$^+$ area coverage and density. For graphs in **(a)**, **(d)**, **(f)**, **(g)**, $n$ = 4 control and 5 cKO mice, unpaired two-tailed $t$ test. Data are shown as mean ± SEM.
